# Structural insights into SETD3-mediated histidine methylation on β-actin

**DOI:** 10.1101/476705

**Authors:** Qiong Guo, Shanhui Liao, Sebastian Kwiatkowski, Weronika Tomaka, Huijuan Yu, Gao Wu, Xiaoming Tu, Jinrong Min, Jakub Drozak, Chao Xu

**Author notes:** These authors contributed equally to this work (Qiong Guo, Shanhui Liao, Sebastian Kwiatkowski). To whom correspondence should be addressed: Chao Xu,; Jakub Drozak,; Shanhui Liao,.

## Abstract

SETD3 is a member of SET (Su(var)3-9, Enhancer of zeste, and Trithorax) domain protein superfamily and plays important roles in hypoxic pulmonary hypertension, muscle differentiation, and carcinogenesis. In a previous paper (*Kwiatkowski* et al. 2018), we have identified SETD3 as the actin-specific methyltransferase that methylates the N^3^ of His73 on β-actin. Here we present two structures of *S*-adenosyl-L-homocysteine-bound SETD3 in complex with either an unmodified β-actin peptide or its His-methylated variant. Structural analyses supported by the site-directed mutagenesis experiments and the enzyme activity assays indicated that the recognition and methylation of β-actin by SETD3 is highly sequence specific, and both SETD3 and β-actin adopt pronounce conformational changes upon binding to each other. In conclusion, the structural research uncovers the molecular mechanism of sequence-selective histidine methylation by SETD3, which not only throws light on protein histidine methylation phenomenon, but also facilitates the design of small molecule inhibitors of SETD3.

## INTRODUCTION

Microfilaments are the building blocks of cytoskeleton and are made up of actin proteins (dos Remedios et al., 2003; Theriot and Mitchison, 1991). There are six actin isoforms in mammalian cells that are characterized based on different expression profiles and cellular functions, including α_skeletal_-, α_cardiac_-, α_smooth_-, β_cyto_-, γ_cyto_-, and γ_smooth_- actins (Gunning et al., 1983; Herman, 1993; Perrin and Ervasti, 2010). Among them, β-actin is ubiquitously expressed and plays critical roles in a wide variety of cellular functions, such as cytoskeleton formation, cell motility and maintenance of cell stability (Leterrier et al., 2017; Nudel et al., 1983).

Many different types of post-translational modifications (PTMs) have been found in actin protein, including acetylation, methylation, SUMOylation and ubiquitination (Terman and Kashina, 2013). N^3^-methylation of His73 in β-actin has been of special interest since its identification in 1967 (Johnson et al., 1967), which seems to be not only due to an extreme evolutionary conservation of this PTM– methylated His is present in almost all eukaryotic actins, but mainly because the methylation of H73 decreases the hydrolysis rate of the actin-bound ATP (Kabsch et al., 1990), as evidenced by the higher ATP exchange rate in the actin-H73A mutant (Nyman et al., 2002).

Although the presence of actin-specific methyltransferases was shown in rabbit muscles decades ago (Raghavan et al., 1992; Vijayasarathy and Rao, 1987), the molecular identity of this enzyme has long been unknown. Recently, we and others identified SETD3 as the actin-specific histidine *N*-methyltransferase that specifically catalyzes the methylation of actin at His73 (Kwiatkowski et al., 2018; Wilkinson et al., 2019). SETD3 is a member of SET domain family, which is a ∼130 aa motif initially identified in Drosophila proteins, with its acronym derived from three histone lysine methyltransferases, **<u>S</u>**u(var)3-9, **<u>E</u>**nhancer of zeste, and **<u>T</u>**rithorax (Dillon et al., 2005). Thus far, most histone lysine methyltransferases contain a SET domain, however, some exceptions include DOT1L (Min et al., 2003), METTL12 (Rhein et al., 2017), and METTL21B (Malecki et al., 2017), among others. Before SETD3 was identified as an actin-specific histidine methyltransferase, SET domain proteins had been known to act as lysine methyltransferases, transferring a methyl moiety from the methyl donor S-Adenosyl methionine (AdoMet) to the substrates to generate the methylated form, with *S*-adenosyl-L-homocysteine (AdoHcy) as the co-product (Dillon et al., 2005).

Since (i) SETD3 has been previously identified as a histone methyltransferase that methylates histone H3 at Lys4 and Lys36 (Eom et al., 2011; Wagner and Carpenter, 2012) and (ii) the results of our preliminary experiments with the use of isothermal calorimetry titration (ITC) and mass spectrometry revealed that SETD3 binds to and methylates a β-actin peptide containing His73 (66-88), but not H3K4 (1-23) or H3K36 (25-47) peptides, we crystallized and solved two structures of AdoHcy-bound SETD3 in complex with either His73 peptide or 3-methylhistidine (His73me) one to understand the molecular mechanism of actin histidine methylation by SETD3.

With the two solved β-actin peptide-SETD3 structures, we uncover that SETD3 recognizes a fragment of β-actin in a sequence-dependent manner and utilizes a specific pocket to catalyze the N^3^-methylation of His73. Moreover, a comprehensive structural, biochemical and enzymatic profiling of SETD3 allows us to pinpoint its key residues important for substrate recognition and subsequent methylation. Therefore, the structural research, supplemented by biochemical and enzymatic experiments, not only provides insights into the catalytic mechanism of SETD3, but also will facilitate the design of specific inhibitors of SETD3 enzyme.

## RESULTS

### SETD3 binds to and methylates β-actin

Since SETD3 was identified as a histidine methyltransferase that methylates His73 of β-actin (Kwiatkowski et al., 2018, Wilkinson et al., 2019), we purified the core region of SETD3 (aa 2-502) and studied by ITC its binding to a His73-containing fragment of β-actin (aa 66-88) (Figures 1A). The ITC binding experiment showed that SETD3 bound to the β-actin peptide with a Kd of 0.17 μM (Figure 1B and Table 1). Given that SETD3 was also reported to be a putative lysine methyltransferase that methylates Lys4 and Lys36 of histone H3 (Eom et al., 2011), we also verified the binding of SETD3 to two different histone peptides, H3K4(1-23) and H3K36(25-47), and found that neither of them binds to SETD3 (Table 1).

**Figure 1.**
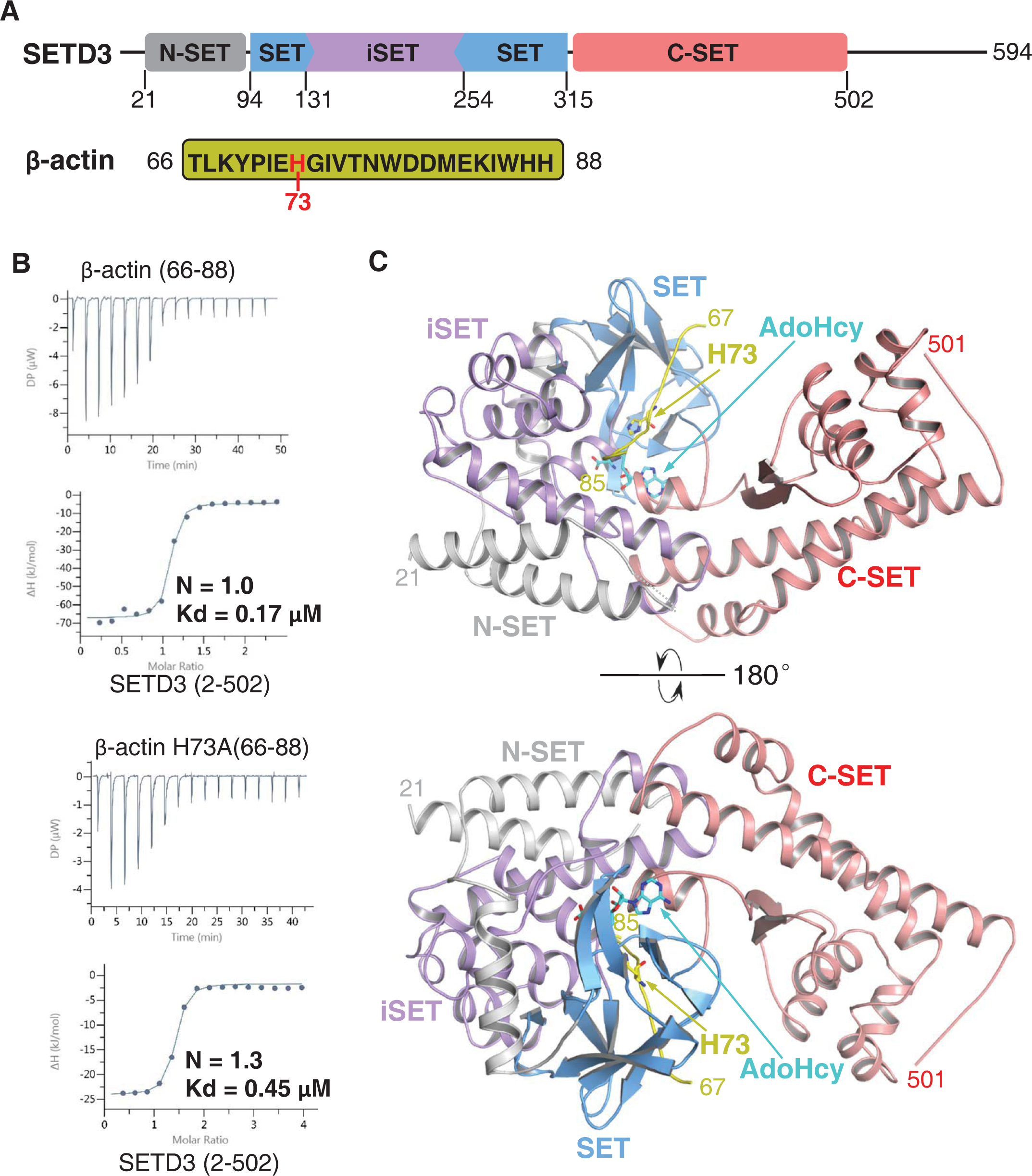
SETD3 core region specifically recognizes a fragment of β-actin containing His73. (A) Domain architecture of full-length human SETD3 (aa 1-594) and the sequence of the β-actin peptide (66-88), with His73 of β-actin highlighted. (B) Representative ITC binding curves for the binding of the SETD3 (aa 2-502) to β-actin peptides of different lengths. Molecular ratios, derived Kds and respective standard deviations are also indicated. (C) Overall structure of AdoHcy bound SETD3 with unmodified β-actin peptide. The SETD3 are colored in the same mode as shown in Figure 1a, with the N-SET, SET, iSET and C-SET regions of SETD3 colored in grey, blue, purple and salmon, respectively. The peptide are shown in yellow cartoon, while His73 and AdoHcy are shown in yellow and cyan sticks, respectively. His73 of actin and the AdoHcy are labeled.

**Table 1.**
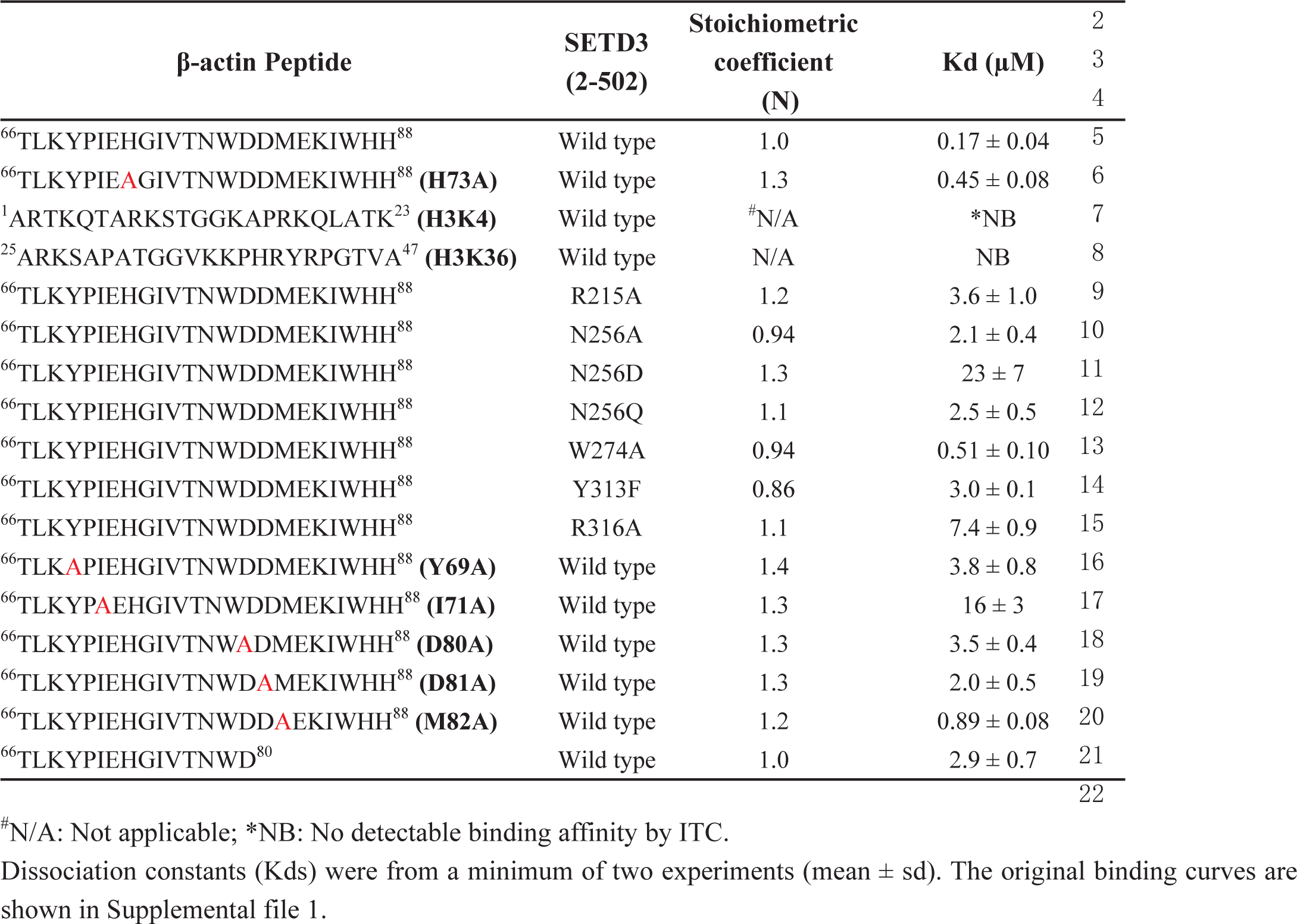
Binding affinities of SETD3 proteins to β-actin peptides

Furthermore, we tested the activity of SETD3 on β-actin(66-88), H3K4(1-23), and H3K36(25-47) by mass spectrometry, and we found that SETD3 methylates β-actin peptide (Figure 1–figure supplement 1A), but does not modify either H3K4 or H3K36 (Figure 1–figure supplement 1B-1C). No methylated product was detected for any of above peptides in the presence of AdoMet without the addition of SETD3 (Figure 1–figure supplement 1A-1C). Moreover, SETD3 did not methylate β-actin(66-88) H73A mutant, although the mutant peptide bound to SETD3 only ∼2.5-fold weaker than the wild type one (0.45 μM vs. 0.17 μM) (Figure 1B and Figure 1–figure supplement 1D). Collectively, the peptide binding experiments and mass spectrometry data indicate that SETD3 specifically binds to the β-actin peptide and methylates it at His73.

### Overall structure of SETD3

To uncover the mechanisms underlying recognition and methylation of β-actin by SETD3, we attempted to crystalize AdoMet-bound SETD3 with full length β-actin, but failed to obtain diffractable crystals. By using the core region of SETD3(2-502) with the β-actin peptide (66-88), we succeeded in the crystallization of the complex and obtained a 1.95 Å structure of AdoHcy-bound SETD3 (2-502) with β-actin (66-88) (Table 2, Figure 1–figure supplement 2A, Figure 1–figure supplement 3). There are four molecules in a crystallographic asymmetric unit and residues 22-501 of SETD3 and 66-84 of β-actin are visible in the complex structures (Figure 1–figure supplement 4A). Based on the density map, we found that His73 of β-actin had been methylated in the structure (Figure 1–figure supplement 5A), although we used an unmethylated peptide. The reason could be that SETD3 bound to the methyl donor AdoMet from *E. coli*, which we used to express our protein, and that the AdoMet-bound SETD3 methylated the peptide at His73 during crystallization. Therefore, this complex represents a snapshot of the post-methyl transfer state, which prompted us to crystallize the substrate-bound enzyme.

**Figure 2.**
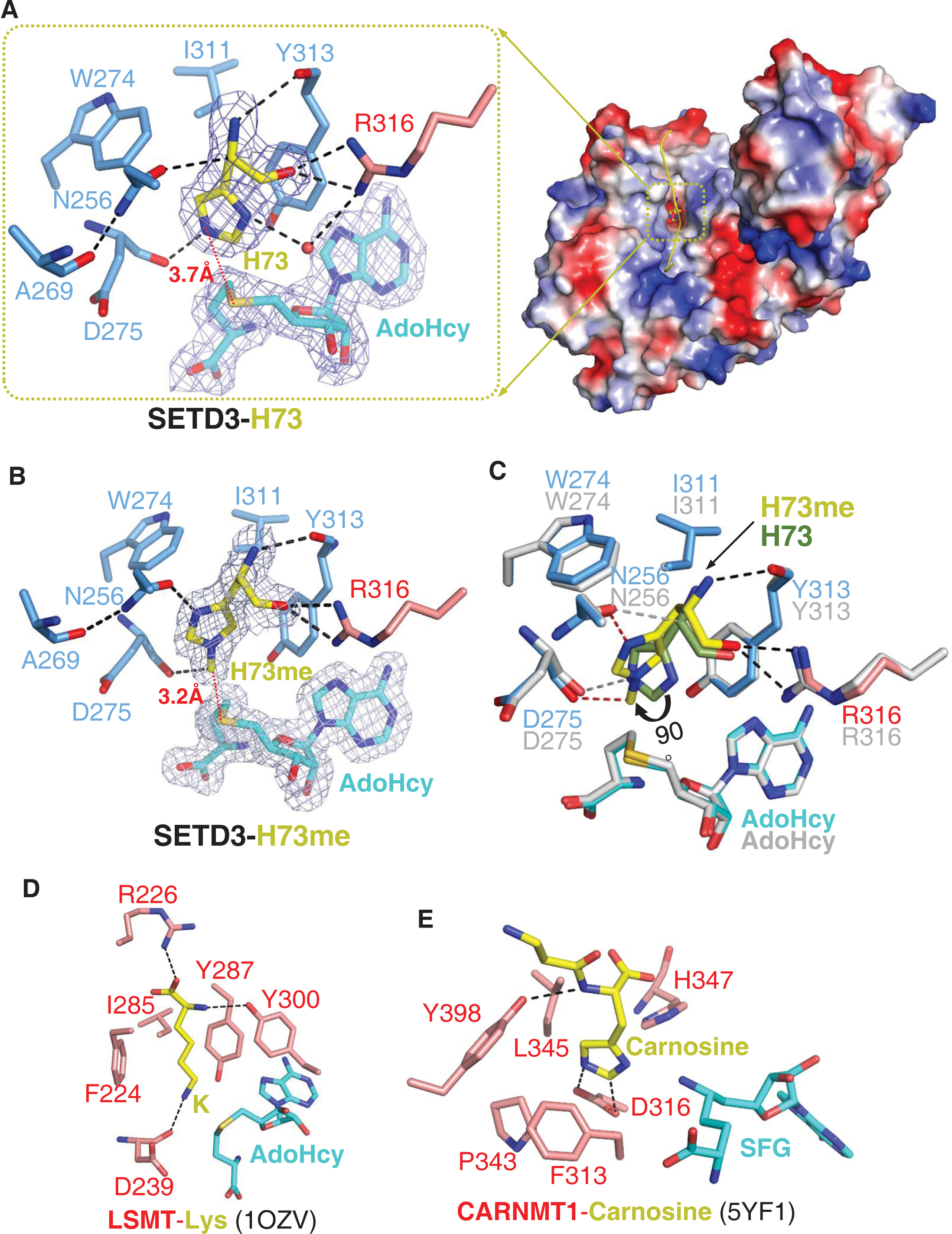
Molecular mechanism of substrate recognition and histidine methylation by SETD3. (A) The β-actin peptide is bound in a long groove on the surface of SETD3 N-lobe region (right), with His73 of β-actin positioned into a hydrophobic pocket (left). His73 and AdoHcy are shown in yellow and cyan sticks, respectively, and their 2|Fo|–|Fc| σ-weighted maps are contoured at 1.2 σ. His73 binding residues of SETD3 are colored based on the regions that they reside as shown in Figure 1A. (B) Detailed interactions between His73me and SETD3 in the post-methyl transfer complex. His73me and AdoHcy are shown in yellow and cyan sticks, respectively, and their 2|Fo|–|Fc| σ-weighted maps are contoured at 1.2 σ. His73me binding residues of SETD3 are colored in the same mode as shown in Figure 2A. (C) Superposition of the two complexes on the histidine/methylhistidine binding pocket. For the SETD3-His73me complex, His73me binding residues of SETD3 are shown in the same way as in Figure 2A, with His73me and AdoHcy shown in yellow and cyan sticks, respectively. For the SETD3-His73 complex, His73 binding residues and AdoHcy are shown in grey sticks, while His73 are shown in green sticks to show that it rotates 90 degree after catalysis. After methylation, one new hydrogen bond is formed between Asn256 and N^1^ atom of His73me (red dash). (D) lysine recognition by LSMT(PDB id: 1OZV) in the presence of AdoHcy. (E) carnosine recognition by CARNMT1 (PDB id: 5YF1) in the presence of SFG. Histidine, lysine and carnosine are shown in yellow sticks, while the protein residues involved in binding are shown in red sticks. AdoHcy and its mimic, SFG, are shown in cyan sticks.

**Figure 3.**
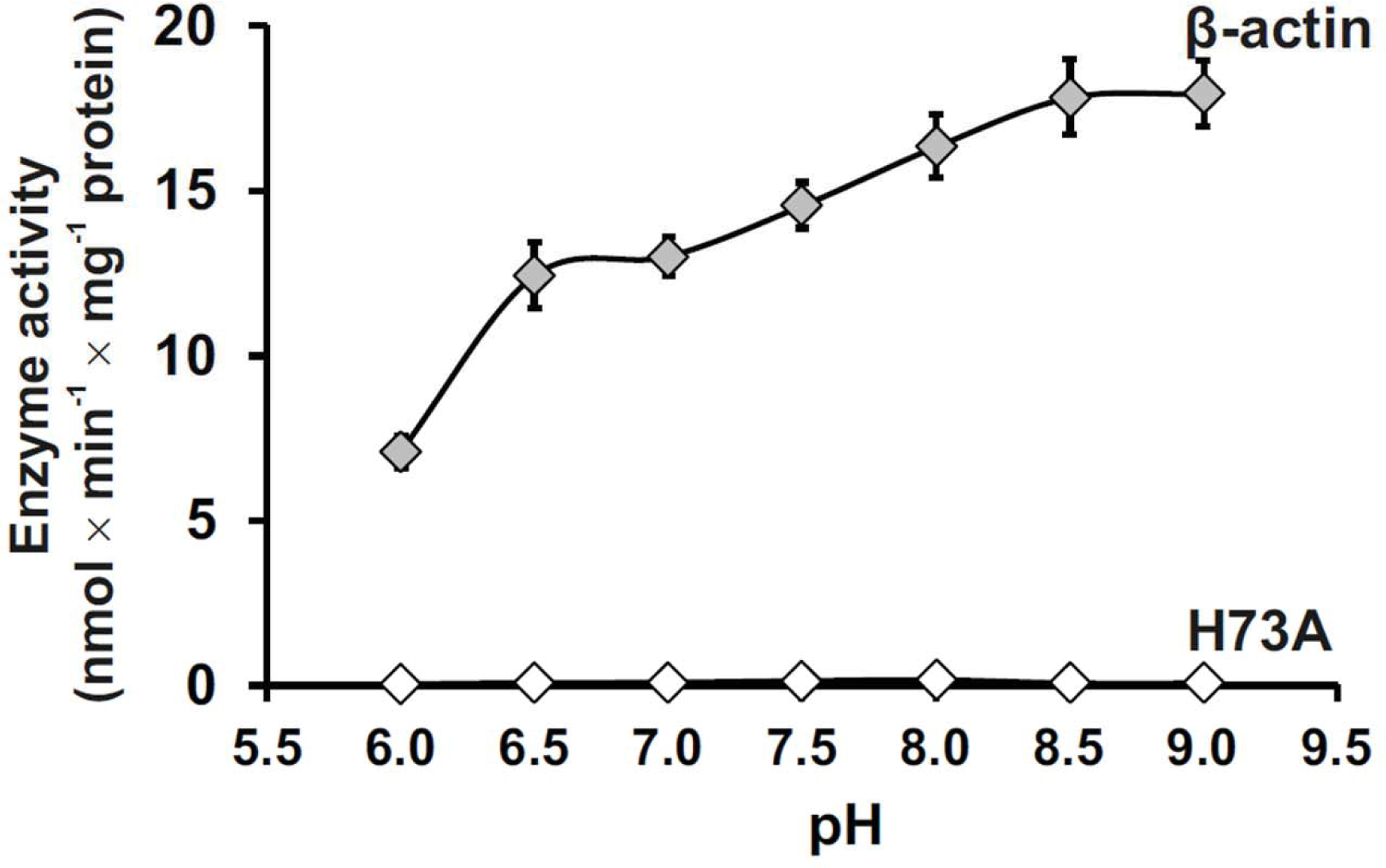
Effect of pH on SETD3 enzyme activity. pH dependence of human SETD3 was determined with the use of purified recombinant N-terminal His6-tagged SETD3 protein (2-502). The enzyme preparation (0.045 μg protein) was incubated at 37°C for 10 min in the presence of 0.8 μM [^1^H+^3^H] AdoMet (80 pmol, ≈350 × 10^3^ cpm) and either 5 μM (500 pmol, 22.34 μg) purified recombinant human β-actin or its mutated form (H73A) which served as a negative control. In all experiments, the reaction mixture contained the homogenous recombinant AdoHcy nucleosidase (1.6 mg protein, 600 nM, E. coli) and adenine deaminase (3.9 mg protein, 600 nM, B. subtilis) to prevent AdoHcy accumulation. The reaction was stopped and the proteins present in the assay mixture were precipitated by adding 10% trichloroacetic acid. This allowed for the separation and specific measurement of the radioactivity incorporated in the protein pellet (extent of actin methylation) from the total radioactivity present in the assay mixture. Values are the means ± S.E. (error bars) of three separate experiments. If no an error bar is visible, it is shorter than the size of the symbol.

**Table 2.** Data collection and refinement statistics

| | SETD3-β-actin | SETD3-methylated β-actin |
| --- | --- | --- |
| **PDB id** | 6ICV | 6ICT |
| **Data collection** |  |  |
| Radiation wavelength (Å) | 0.9789 | 0.9792 |
| Space group | P 1 21 1 | P 1 21 1 |
| Cell dimensions |  |  |
| *a*, *b*, *c* (Å) | 60.19, 175.17, 66.50 | 59.98,176.69,125.89 |
| $\alpha$, $\beta$, $\gamma$ (°) | 90, 92.57, 90 | 90,93.37,90 |
| Resolution (Å) | 35.00-2.15(2.23-2.15) | 72.27-1.95(2.06-1.95) |
| R<sub>merge</sub> | 0.135(0.609) | 0.169(0.868) |
| *I* / σ*I* | 14(3.4) | 8.1(2.6) |
| CC1/2 | 0.988(0.884) | 0.991(0.827) |
| Completeness (%) | 100(100) | 99.1(99.7) |
| Redundancy | 6.4(6.4) | 7.0(7.0) |
| **Refinement** |  |  |
| Resolution (Å) | 35.00-2.15 | 59.87-1.95 |
| No. of reflections (used/free) | 71998/3629 | 187163/9084 |
| R<sub>work</sub>/R<sub>free</sub> | 0.168/0.205 | 0.178/0.214 |
| Number of atoms/B-factor (Å<sup>2</sup>) | 8493/30.7 | 16956/33.7 |
| Protein | 7574/30.3 | 15147/33.1 |
| Peptide | 276/32.9 | 591/38.4 |
| AdoHcy | 52/19.3 | 104/20.0 |
| Solvent | 591/36.2 | 1114/40.0 |
| RMSD bonds (Å)/angles (°) | 0.007/0.84 | 0.007/0.87 |
| Ramachandran Plot |  |  |
| favored/allowed/outliers (%) | 98.76/1.24/0 | 98.98/1.02/0 |

Values in parentheses are for highest-resolution shell.

To avoid the methylation reaction, SETD3 was purified by adding 5-fold excess of AdoHcy in the buffer to compete off AdoMet. Then the non-methylated peptide was mixed with the purified SETD3 to form the complex. The complex was crystallized and solved at a resolution of 2.15 Å (Table 2). There are two molecules in a crystallographic asymmetric unit (Figure 1–figure supplement 4B). In this structure, the density map of the peptide indicates that it was unmodified, suggesting that methylation did not occur (Figure 1–figure supplement 5B). The complex structure hereby reflects the image of the pre-methyl transfer state.

In both structures, the SETD3 core region adopts a V-shape architecture and is composed of N-SET (N-terminal to the SET domain), SET, iSET (an insertion in the SET domain), and C-SET (C-terminal to the SET domain) domains (Figure 1C). The N-terminal lobe consists of the N-SET (α1-α3), SET (β1-β12), and iSET (α4-α11) domains (Figure 1–figure supplement 2A, Figure 1–figure supplement 3). The N-SET domain consists of three α helices (α1-α3), and encompasses the SET and iSET domains (Figure 1–figure supplement 2A, Figure 1–figure supplement 3). While α1 and α2 of N-SET form a 4-helix bundle with α10-α11 of iSET, α3 of N-SET is localized at the SET-iSET interface and comes into contact with the anti-parallel β strands (β1-β2) of SET and α8 of iSET (Figure 1C, Figure 1–figure supplement 3). The SET domain adopts a canonical SET domain fold similar to that of LSMT (Trievel et al., 2002), with twelve β strands arranged into four anti-parallel β sheets (β1-β2, β3-β11, β4-β10-β9, β5-β7-β6) and one parallel β sheet (β8-β12) (Figure 1–figure supplement 2A, Figure 1–figure supplement 3). The iSET domain is a helical region (α4-α11) inserted between β5 and β6 of the SET domain (Figure 1–figure supplement 2A, Figure 1–figure supplement 3). The C-terminal lobe, only consisting of the C-SET domain, is almost entirely helical (α12-α19) except two anti-parallel strands (β13-β14), with α12 and the N-terminal end of α19 packed against α10 and α9 of iSET, respectively (Figure 1–figure supplement 2A, Figure 1–figure supplement 3). Consistently, a database search using the DALI server (Holm and Rosenstrom, 2010) revealed that the overall fold of SETD3 is highly similar to those of LSMT (Z-core: 31.4, RMSD: 3.8 Å for 425 Cα atoms) and SETD6 (Z-core: 28.7, RMSD: 2.8 Å for 421 Cα atoms) despite the low sequence identities (24%-25%).

### AdoHcy binding pocket within SETD3

In both complex structures, SETD3 binds AdoHcy in a cleft formed between SET and C-SET and buttressed by iSET at the bottom (Figure 1C). Specifically, the adenine ring of AdoHcy is sandwiched between the side chain of Glu104 and the aromatic ring of Phe327, forming stacking and π-π interactions, respectively (Figure 1–figure supplement 6A). The AdoHcy N^6^ and N^7^ atoms are hydrogen bonded to the main chain carbonyl and amide groups of His279, respectively, while the AdoHcy C^8^ atom forms one C-H…O hydrogen bond with the hydroxyl group of Tyr313 (Figure 1–figure supplement 6A).

In addition to the adenine specific interactions, the ribose O3’ and O4’ atoms of AdoHcy form hydrogen bonds with the main chain carbonyl of Ser325 and the side chain carboxyl of Asn278, respectively (Figure 1–figure supplement 6A). The side chain carboxyl of Asn278 makes another hydrogen bond with the AdoHcy amide, while the latter is also H-bonded to the main chain carbonyl of Phe106 (Figure 1–figure supplement 6A). The AdoHcy carboxylate forms four hydrogen bonds with SETD3 residues, one with the main chain amide of Phe106, one with the guanidino group of Arg254, and the other two with the guanidino group of Arg75 (Figure 1–figure supplement 6A). All AdoHcy binding residues are conserved in other SETD3 orthologs, suggesting a conserved function of SETD3 (Figure 1–figure supplement 2A).

The AdoHcy binding mode of SETD3 is similar to those observed in some other SET domain structures, such as those of LSMT (Trievel et al., 2002) and SETD6 (Chang et al., 2011) (Figure 1–figure supplement 6B-6C). Among these AdoHcy binding residues, Asn278, His279 and Phe327 are conserved in LSMT and SETD6, while Glu104 and Arg254 are conserved only in LSMT, but not in SETD6 (Figure 1–figure supplement 2B, Figure 1–figure supplement 6B-6C). Arg75 is an AdoHcy binding residue only found in SETD3 (Figure 1–figure supplement 2B).

### β-actin binding mode of SETD3

In both complexes, the β-actin peptide lies in a narrow groove formed by SET, iSET and C-SET, with the unmodified or methylated His73 residue accommodated in a pocket composed of α11 of iSET, β6 and β12 of SET, and α12 of C-SET, and further buttressed by η5 of SET (Figure 1C, Figure 2A). When superimposing our substrate-bound complex structure with the previously determined AdoMet-SETD3 binary structure (PDB id: 3SMT) on SETD3, we found that, despite the overall structural similarity between two SETD3 structures (with a RMSD of 0.66 Å in Cα positions), the region containing two β-sheets of the SET domain (β4-β10-β9 and β3-β11) takes a conformational change upon peptide binding, due to its direct interactions with the N-terminal end of the peptide, with the three loops preceding β4, β9 and β11 shift 4.2 Å, 4.4 Å and 6.8 Å towards the peptide, respectively (Figure 1–figure supplement 7). The detailed interactions between SETD3 and the N-terminal end of the peptide will be described in the following section.

### A unique histidine recognition pocket in SETD3 confers to its histidine methyltransferase activity

In the substrate complex, residues Leu67-Glu83 of the peptide are visible, with His73 occupying a hydrophobic pocket (Figure 2A). The main chain of His73 forms several hydrogen bonds with SETD3, with its main chain amide, Cα and main chain carbonyl H-bonded to the main chain carbonyl of Tyr313, the side chain carboxyl of Asn256, and the guanidino of Arg316, respectively (Figure 2A). The imidazole ring of His73 is parallel to the aromatic ring of Tyr313, with its orientation determined by two hydrogen bonds, one between the N^1^ atom of the ring and the guanidino of Arg316 mediated by a water molecule, and the other between the N^3^ atom of the ring and the main chain carbonyl of Asp275 (Figure 2A). In addition to the stacking interactions with Tyr313, His73 also makes hydrophobic contacts with Trp274 and Ile311. The distance between the N^3^ atom of His73 and the sulfur atom of the AdoHcy is 3.7 Å, suggesting that it represents well the pre-methyl transfer state (Figure 2A).

In the product complex, the residues Leu67-Lys84 and the main chain of Ile85 of the peptide are visible, with His73 methylated to His73me (Figure 1–figure supplement 8). Despite the fact that the methylated peptide adopts a 3^10^ helix at its C-terminal end, the two complexes are highly similar, with a RMSD of 0.19 Å over protein Cα atoms and a RMSD of 0.32 Å over peptide Cα atoms. In the product complex, His73me is inserted into the same pocket of SETD3 as shown in the substrate complex and the distance between the installed methyl group of His73 and the sulfur atom of the AdoHcy is 3.2 Å, with its density map distinguishable from that of unmodified histidine (Figure 2B).

The superposition of the two complexes clearly indicates that the hydrogen bonds between the main chain of His73me and the main chain carbonyl of Tyr313 and the guanidino of Arg316 are intact after methylation (Figure 2B). However, the imidazole ring of His73me is rotated by ∼90° compared to that of the unmodified His73, with its C^4^H pointing toward the ring of Tyr313 (Figures 2B-2C) and its N^1^ atom hydrogen bonded to the side chain carboxyl group of Asn256 (Figure 2B).

When comparing SETD3 with the other SET domain containing lysine methyltransferases, we found that there are two main reasons that SETD3 is not likely to be a lysine methyltransferase. Firstly, the histidine binding pocket of SETD3 is too shallow to accommodate the aliphatic side chain of a lysine. Replacement of His73 with a lysine, would not only disrupt the water mediated hydrogen bond between N^1^ of His73 and Arg316, but could also lead to the steric clash between the aliphatic side chain of lysine and the main chain of the bottom carbonyl cage residue, Asp275. Secondly, the phenylalanine of LSMT and SETD6 that makes cation-π interaction with the lysine, is replaced by an asparagine (Asn256) in SETD3, and Asn256 is critical for substrate binding and histidine methylation (Figures 2A-2B and Figure 1–figure supplement 2A). Therefore, this substitution establishes the imidazole ring-specific interaction with histidine by impairing the lysine specific interaction.

### SETD3 exhibits extensive interactions with β-actin

In both complexes, peptide residues flanking His73 or His73me also interact extensively with the N-terminal lobe of SETD3, with Leu67-Glu72 and Gly74-Glu83 mainly contacting the SET and iSET motifs, respectively (Figure 2–figure supplement 1, Figure 2–figure supplement 2). Leu67 and Pro70 of β-actin make hydrophobic contacts with Ile284 of SETD3 and there are additional hydrophobic interactions between Tyr69 of β-actin and Pro259, Tyr288 and Leu290 of SETD3 (Figure 2–figure supplement 1, Figure 2–figure supplement 2). Two main chain hydrogen bonds are formed between Tyr69 of β-actin and Tyr288 of SETD3, and between Ile71 of β-actin and Thr286 of SETD3, which induces the shift of the two sheets (β4-β10-β9 and β3-β11) in SETD3 as mentioned above (Figure 1–figure supplement 7). Ile71 of actin also makes hydrophobic interactions with Ile258, Ile271, Trp274, Thr286, Tyr288 and Cys295 of SETD3 (Figure 2–figure supplement 1, Figure 2–figure supplement 2). Of note, Leu283-Thr285 of SETD3 adopts a β strand in the structure of AdoMet bound SETD3 without peptide (PDB: 3SMT), but decomposes upon binding to the N-terminal side of β-actin, which shortens the β9 strand of SETD3 in the peptide-bound structure (Figure 1–figure supplement 7). The side chain carboxylate group of Glu72 forms one hydrogen bond with the guanidino group of Arg316, which is the only C-SET domain residue that contacts the N-terminal side of the β-actin peptide (Figure 2–figure supplement 1, Figure 2–figure supplement 2).

Regarding the residues C-terminal to His73 of β-actin, the main chain amide group of Gly74 is hydrogen bonded to the main chain carbonyl group of Gln255, while the side chain of Val76 stacks with the imidazole ring of His324 (Figure 2–figure supplement 1, Figure 2–figure supplement 2). In contrast to Gly74 and Val76, the Ile75 side chain is solvent-exposed and is thereby not involved in the interactions with SETD3 (Figure 2–figure supplement 1). Thr77 not only makes hydrophobic contact with Leu268, but also forms two hydrogen bonds through its hydroxyl group with the side chain amide groups of Asn154 and Gln255, respectively (Figure 2–figure supplement 1, Figure 2–figure supplement 2). The side chain amide group of Asn154 makes another hydrogen bond with the main chain carbonyl of Asn78. The indole ring of Trp79 is inserted into an open hydrophobic pocket composed of Ile155, Val248, Val251 and Met252, with its main chain carbonyl H-bonded to the side chain amide of Gln216, while the carboxyl group of Asp80 is hydrogen bonded to the side chain amide of Arg215 (Figure 2–figure supplement 1, Figure 2–figure supplement 2). Asp81 of β-actin forms three hydrogen bonds with SETD3, one with the side chain amide of Gln216 and two with the guanidino group of Arg215 (Figure 2–figure supplement 1, Figure 2–figure supplement 2). Of note, the peptide sequence is also conserved in α- and γ-actin proteins, suggesting that SETD3 might also recognize and methylate other actin isotypes.

### Key residues of SETD3 involved in β-actin binding and methylation

To evaluate the roles of the SETD3 residues around the active site in β-actin methylation, we made several single mutants and compared quantitatively their binding affinities to the β-actin peptide by ITC. While all SETD3 mutants displayed weaker binding affinities towards the peptide, R215A and R316A exhibited the most significantly reduced binding affinities by 21- and 42- fold, respectively (Table 1). Our complex structures showed that both Arg215 and Arg316 make several hydrogen bonds with β-actin (Figure 2B, Figure 2–figure supplement 1), implicating that they play a critical role in substrate recognition (Table 1). N256A also reduces the β-actin binding affinity by 12-fold (Kd = 2.1 μM) (Table 1), which disrupts not only the hydrogen bond to the main chain of the His73 Cα atom (Figure 2A), but also the one to the N^1^ atom of His73me (Figure 2B). In the substrate complex, the imidazole ring of His73 stacks with the aromatic ring of Tyr313, with the N^3^ atom of His73 in proximity to the hydroxyl group of Tyr313 (Figure 2A). The hydroxyl group of Tyr313 partially neutralizes the charge of N^3^, as verified by the fact that mutating Tyr313 to phenylalanine diminishes binding affinity by 17-fold (Kd = 3.0 μM) (Table 1). A similar favorable charge-charge interaction was also observed in the structure of Gemin5 bound to m^7^G base (Xu et al., 2016),

Given that the β-actin peptide interact with SETD3 via flanking sequences of His73, we performed mutagenesis and ITC studies to corroborate the roles of Tyr69, Ile71, Asp80, Asp81, and Met82. Tyr69, Ile71 and Met82 of β-actin contact SETD3 via hydrophobic interactions, and replacing either of them by alanine dramatically reduces the SETD3 binding affinity by 5-90 fold (Table 1). Consistent with the fact that both Asp80 and Asp81 of β-actin form hydrogen bonds with SETD3, D80A and D81A impair the binding to SETD3 by 20- and 11-fold, respectively (Table 1). The interactions of Asp81 and Met82 of β-actin with SETD3 also explains why truncation of Asp81-His88 of the peptide reduces the binding to SETD3 by ∼17-fold (Figure 2–figure supplement 1, Table 1). Taken together, the flanking sequences of His73, rather than His73 itself on the β-actin peptide, are critical for binding to SETD3, suggesting that the substrate recognition by SETD3 is highly sequence-selective.

To gain further insights into how substrate recognition by SETD3 affects subsequent histidine methylation, we used the mass spectrometry to compare the activity of the above mentioned SETD3 mutants with that of the wild type protein in the presence of AdoMet. The mass spectrometry data indicate that the peaks of products generated by the four SETD3 mutants (R215A, N256A, Y313F, and R316A) were much lower than that observed for wild type SETD3 (Figure 1–figure supplement 1E), suggesting that these mutants displayed weaker histidine methylation activity.

### Activity of SETD3 enzyme and its mutant variants towards β-actin protein

To verify the identification of key catalytic residues of SETD3, and to analyze the roles of substrate recognition residues in catalysis, we determined the kinetic properties of the wild-type and mutant variants of purified recombinant SETD3 in the presence of homogenous full length recombinant human β-actin. All enzyme assays were performed in the presence of *E. coli* AdoHcy nucleosidase and *B. subtilis* adenine deaminase to prevent the accumulation of AdoHcy in the reaction mixture (Dorgan et al., 2006). As shown in Table 3, the wild-type SETD3 has a very high affinity towards both AdoMet (K_M_ ≈ 0.1 μM) and β-actin (K_M_ ≈ 0.5 μM), though it appeared as a sluggish catalyst, with k_cat_ equal to about 0.7-0.8 min^-1^. All mutant enzymes exhibited a reduced catalytic efficiency as indicated by k_cat_/K_M_, 2-400 fold and 1.6-2000 fold for β-acint and AdoMet, respectively (Table 1), confirming the importance of examined amino acid residues for the efficient catalysis.

**Table 3.**
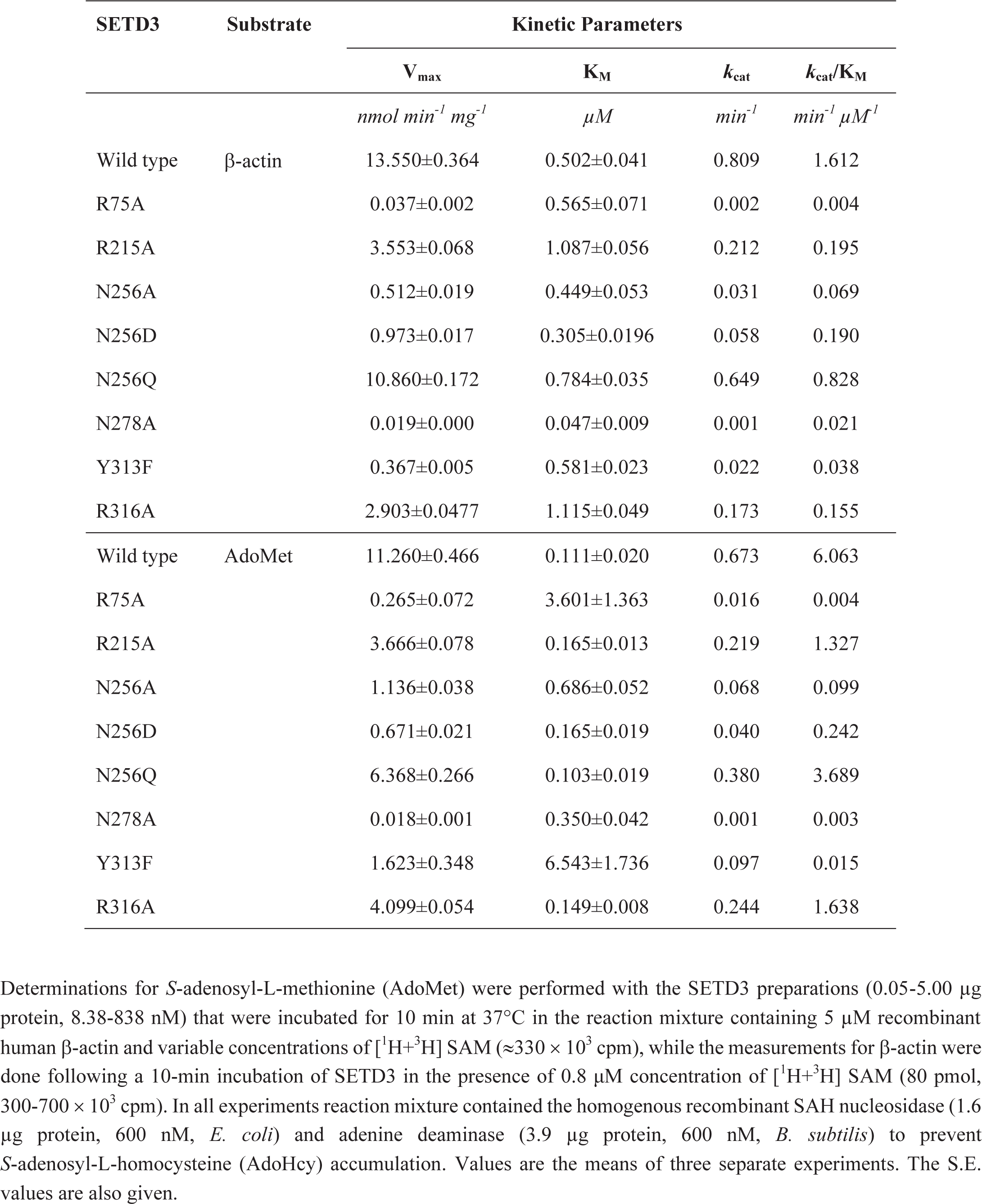
Kinetic properties of wild type SETD3 and its mutants. Kinetic parameters were determined with the use of purified recombinant N-terminal His_6_-tagged SETD3 proteins.

| SETD3 | Substrate | Kinetic Parameters |  |  |  |
| --- | --- | --- | --- | --- | --- |
|  |  | V<sub>max</sub> | K<sub>M</sub> | k<sub>cat</sub> | k<sub>cat</sub>/K<sub>M</sub> |
|  |  | *nmol min<sup>-1</sup> mg<sup>-1</sup>* | *μM* | *min<sup>-1</sup>* | *min<sup>-1</sup> μM<sup>-1</sup>* |
| Wild type | β-actin | 13.550±0.364 | 0.502±0.041 | 0.809 | 1.612 |
| R75A |  | 0.037±0.002 | 0.565±0.071 | 0.002 | 0.004 |
| R215A |  | 3.553±0.068 | 1.087±0.056 | 0.212 | 0.195 |
| N256A |  | 0.512±0.019 | 0.449±0.053 | 0.031 | 0.069 |
| N256D |  | 0.973±0.017 | 0.305±0.0196 | 0.058 | 0.190 |
| N256Q |  | 10.860±0.172 | 0.784±0.035 | 0.649 | 0.828 |
| N278A |  | 0.019±0.000 | 0.047±0.009 | 0.001 | 0.021 |
| Y313F |  | 0.367±0.005 | 0.581±0.023 | 0.022 | 0.038 |
| R316A |  | 2.903±0.0477 | 1.115±0.049 | 0.173 | 0.155 |
| Wild type | AdoMet | 11.260±0.466 | 0.111±0.020 | 0.673 | 6.063 |
| R75A |  | 0.265±0.072 | 3.601±1.363 | 0.016 | 0.004 |
| R215A |  | 3.666±0.078 | 0.165±0.013 | 0.219 | 1.327 |
| N256A |  | 1.136±0.038 | 0.686±0.052 | 0.068 | 0.099 |
| N256D |  | 0.671±0.021 | 0.165±0.019 | 0.040 | 0.242 |
| N256Q |  | 6.368±0.266 | 0.103±0.019 | 0.380 | 3.689 |
| N278A |  | 0.018±0.001 | 0.350±0.042 | 0.001 | 0.003 |
| Y313F |  | 1.623±0.348 | 6.543±1.736 | 0.097 | 0.015 |
| R316A |  | 4.099±0.054 | 0.149±0.008 | 0.244 | 1.638 |

Determinations for *S*-adenosyl-L-methionine (AdoMet) were performed with the SETD3 preparations (0.05-5.00 μg protein, 8.38-838 nM) that were incubated for 10 min at 37°C in the reaction mixture containing 5 μM recombinant human β-actin and variable concentrations of [<sup>1</sup>H+<sup>3</sup>H] SAM (≈330 × 10<sup>3</sup> cpm), while the measurements for β-actin were done following a 10-min incubation of SETD3 in the presence of 0.8 μM concentration of [<sup>1</sup>H+<sup>3</sup>H] SAM (80 pmol, 300-700 × 10<sup>3</sup> cpm). In all experiments reaction mixture contained the homogenous recombinant SAH nucleosidase (1.6 μg protein, 600 nM, *E. coli*) and adenine deaminase (3.9 μg protein, 600 nM, *B. subtilis*) to prevent *S*-adenosyl-L-homocysteine (AdoHcy) accumulation. Values are the means of three separate experiments. The S.E. values are also given.

Arg75 and Asn278 are both AdoMet binding residues and mutating either of them to alanine severely impaired k_cat_ of the enzyme towards β-actin and AdoMet (c.f. Table 3). Interestingly, although N256A and Y313F weakened the binding to β-actin peptide by ∼12- and ∼17- fold, respectively (Table 1), both mutants displayed K_M_ values (0.449 μ*M* and 0.581 μ*M*, respectively) comparable to that of wild type SETD3 (0.502 μ*M*), probably owning to their much lower k_cat_ values (∼26- and ∼36-fold, respectively) than wild type SETD3 (c.f. Table 3). For AdoMet, N256A and Y313F both have a higher K_M_ (∼6- and ∼58-fold, respectively) and lower k_cat_ (∼9- and ∼6-fold, respectively) than wild type enzyme. We also examined the kinetic properties of N256D and N256Q, and found that for both β-actin and AdoMet, kinetic parameters of N256Q are comparable to those of the wild type SETD3, while N256D behaves similarly to N256A (Table 3). Taking into account that Asn256 and Tyr313 of SETD3 interact mainly with His73 of β-actin, and not with AdoMet (Figures 2A-2B), these results may imply that Asn256 and Tyr313 of SETD3 plausibly bind to His73 of β-actin by favorably orienting its imidazole ring to facilitate the efficient transfer of the methyl group from AdoMet to His73. The consequence of the N256A and Y313F mutations is thus a decrease in the rate of “successful” methylation events that could be partially overcome by an increase in AdoMet concentration.

In contrast to Asn256 and Tyr313, Arg215 and Arg316 of SETD3 contact flanking sequences of His73. R215A and R316A, which impaired the binding of SETD3 to β-actin peptide, also displayed higher K_M_ values (∼2 fold for both) and lower k_cat_ values (∼3.5- and ∼4.5-fold, respectively) for β-actin protein (Table 3), confirming that those SETD3 residues involved in β-actin peptide recognition are also important for efficient His73 methylation in the context of the full length β-actin.

We further investigated the pH-dependent activity of SETD3 towards full length β-actin and found that SETD3 catalyzes the methylation of β-actin in a wide range of pH (6.0-9.0), with its activity increasing with pH and reaching maximum at pH 9.0 (Figure 3). The detectable activity of SETD3 at neutral pH is consistent with the fact that histidine deprotonation occurs at neutral pH, which facilitates the transfer of methyl group from AdoMet to histidine. Like lysine, histidine deprotonation also increases with pH, well explaining the higher activity of SETD3 in alkaline conditions.

## DISCUSSION

### Catalytic mechanism of SETD3

As the major component of microfilament, actin participates in diverse cellular processes. Different types of PTMs have been identified in actin, which play important roles in mediating actin functions *in vivo*. (Terman and Kashina, 2013). N^3^-methylhistidine was identified in β-actin decades ago (Johnson et al., 1967) and was postulated to mediate the polymerization and hydrolysis of actin filaments (Kabsch et al., 1990). However, how the mark is incorporated, remains enigmatic. Our identification of SETD3 as a histidine methyltransferase catalyzing the His73 methylation of β-actin, answers this unsolved question.

Although SETD3 contains a canonical SET domain and its AdoHcy binding mode is similar to those observed in other SET domain lysine methyltransferases, the predicted SET domain fold of SETD3 does not help to disclose the molecular mechanisms of the substrate recognition and histidine methylation, since the SET domain itself, as a short motif of ∼110 aa, always requires iSET and C-SET domains to complete its function (Chang et al., 2011; Trievel et al., 2002). Our solved substrate- and product-bound structures of SETD3, provides an insight into the pre- and post-methyl transfer states of the histidine methylation by the same enzyme. The SETD3 structures, for the first time, uncover the molecular mechanism by which a SET domain protein acts as the actin His73-specific methyltransferase.

Our structural analyses, assisted by mutagenesis and biochemical experiments, suggest that the recognition of β-actin and subsequent catalysis by SETD3 is different from those observed for other SET domain-containing lysine methyltransferases in following aspects: i) the histidine recognition pocket of SETD3 is shallower and key residues of SET3, which form hydrogen bonds with the His73 imidazole ring, are not conserved in other SET proteins. (Figures 2A-2B, Figure 1–figure supplement 2B); ii) In two solved SETD3 complex structures, the methylated histidine rotates its side chain by 90° (Figure 2C); (iii) The recognition of β-actin by SETD3 is highly dependent on the flanking sequences of His73 (Figure 2–figure supplement 1), which is consistent with the high binding affinity between SETD3 and β-actin H73A peptide (66-88) (Figure 1B). Although N^3^-methylhistidine is also found in other proteins, such as mammalian myosin (Johnson et al., 1967) and *Saccharomyces cerevisiae* Rpl3 (Webb et al., 2010), the sequence preference suggests that SETD3 probably works only on His73 of β-actin, as well as on corresponding histidines in α- and γ-actins. One implication is that no SETD3 ortholog is found in baker’s yeast.

Previously, we identified *Carnosine N-methyltransferase 1* (*CARNMT1*) as the gene coding for carnosine *N*-methyltransferase in mammals (Drozak et al., 2015), which catalyzes N^1^-methylation of the histidine imidazole ring of carnosine (13-alanyl-L-histidine), an abundant dipeptide in skeletal muscle of vertebrates. Recently, the structures of CARNMT1 bound to analogs of N^1^-methylhistidine were also reported (Cao et al., 2018), allowing us to compare the N^3^-histidine methylation by SETD3 with the N^1^-histidine methylation by CARNMT1 through comparing two complexes (Figures 2A and 2E). We found that SETD3 and CARNMT1 not only adopt different folds and AdoHcy binding modes, but also display distinct histidine recognition and catalysis mechanisms. Asp316 of CARNMT1 forms two hydrogen bonds with the imidazole ring of the histidine to facilitate its deprotonation (Figure 2E) (Cao et al., 2018). In the SETD3-substrate complex, His73 stacks with Tyr313 and is hydrogen bonded to the main chain carbonyl group of Asp275 (Figure 2A), which orients the His73 imidazole ring for efficient catalysis. Tyr313 of SETD3 plays important roles in substrate recognition and subsequent histidine methylation, supplemented by the mutagenesis, ITC binding experiments and enzymatic assays (Table 1, Table 3). Finally, the 90-degree rotation of the His73me imidazole ring was not found in CARNMT1-mediated histidine methylation.

### Implications of conformational changes of actin upon binding to SETD3

In natively purified rabbit β-actin, His73 exists as the methylated form and it is localized close to the phosphate groups of ATP binding to actin (Kabsch et al., 1990). Although our complexes only contain a fragment of β-actin, they could still provide clues to the role of SETD3 in mediating actin functions. The residues 66-84 of β-actin consist of three β strands followed by an α-helix, however, the β-actin peptide adopts an extended conformation upon binding to SETD3 (Figure 2–figure supplement 3), suggesting that β-actin probably endures local structural remodeling during methylation. Given that SETD3 is found to be co-purified with other factors, such as the peptide release factor ERF3A (Kwiatkowski et al., 2018), SETD3 might require the assistance of other protein(s) to acquire the methylation of actin His73.

### Conclusions

In the current investigation, we have identified molecular mechanism of β-actin methylation by SETD3 and indicated that the enzyme displays a very high substrate specificity towards β-actin His73. Since the overexpression of SETD3 was recently reported to be associated with cancer malignancy (Cheng et al., 2017), the high-resolution complex structures of SETD3 might be helpful for a better understanding of the role of actin histidine methylation in human cancer, as well as they could provide a structural basis for designing specific SETD3 inhibitors in the near future.

During the peer review process of our manuscript, a manuscript was published uncovering the function of mammalian SETD3 in preventing primary dystocia and presenting the structures of SETD3 bound to an unmodified actin peptide (66-80) (Wilkinson et al., 2019). The structural findings reported in both manuscripts are similar except that our complex structures contain a longer actin fragment (66-88) and that the histidine is methylated in one of our complex.

Materials and Methods.

| Reagent type<br>(species) or<br>resource | Designation | Source or<br>reference | Identifiers | Additional information |
| --- | --- | --- | --- | --- |
| gene (*Escherichia coli*) | AdoHcy nucleosidase | NA | NCBI: NC_000913.3 |  |
| gene (*Bacillus subtilis*) | adenine deaminase | NA | NCBI: NC_000964.3 |  |
| gene (*H.sapiens*) | β-actin | NA | GenBank: NM_001101.4 | |
| gene (*H.sapiens*) | SETD3 | NA | NCBI: NM_032233.2 |  |
| recombinant DNA<br>reagent | pCOLD I (plasmid) | Takara Bio | 3361 |  |
| genetic reagent (E.coli) | pCOLD I/β-actin | PMID: 20851184 | | Kind gift of M. Tamura,<br>Ehime University |
| genetic reagent (E.coli) | pCOLD I/SAH<br>nucleosidase | PMID: 30526847 |  | Overexpression of the<br>recombinant *E.coli* AdoHcy<br>nucleosidase in *E.coli* BL21<br>(DE3) |
| genetic reagent (E.coli) | pCOLD I/adenine<br>deaminase | PMID: 30526847 |  | Overexpression of the<br>recombinant *B.subtilis*<br>adenine deaminase in *E.coli*<br>BL21 (DE3) |
| strain, strain<br>background<br>(*B.subtilis*) | *Bacillus subtilis* | Sigma-Aldrich | ATCC:6633 |  |
| strain, strain<br>background (*E.coli*) | *E.coli* BL21 (DE3) | Agilent<br>Technologies | 200131 |  |
| chemical compound,<br>drug | S-[methyl-<sup>3</sup>H]adenosyl<br>-L-methionine;<br>[<sup>3</sup>H]SAM | PerkinElmer | NET155V250UC;<br>NET155V001MC |  |
| software, algorithm | GraphPad Prism | GraphPad<br>Software | RRID:SCR_002798 |  |
| pET28-MHL | vector | Addgene | 26096 (Genbank<br>accession number:<br>EF456735) | Expression of SETD3 and its<br>mutants |
| Origin 6.1, Origin 7.0 | software | OriginLab | http://www.originlab.com/ | For ITC curve fitting and<br>calculation Kd values. |
| HKL2000 | software | PMID: 27754618 | http://www.hkl-xray.com/ | Processing the crystal<br>structure data |
| PHASER | software | PMID: 19461840 | http://www.phaser.cimr.cam.ac.uk/ | Molecular replacement |
| PHENIX | software | PMID: 20124702 | http://www.phenix-online.org/ | Structure refinement |
| Coot | software | PMID: 15572765 | http://www2.mrc-lmb.cam.ac.uk/Personal/pemsley/coot/ | Structure refinement |
| Pymol | software | DeLano Scientific LLC | http://www.pymol.org/ | Making structure figures |
| MOLPROBITY | software | PMID: 20057044 | http://molprobity.biochem.duke.edu/ | Structure validation |

### Cloning, protein expression and purification of SETD3

Gene encoding the core region of SETD3 (2-502) were synthesized by Sangon Biotech (Shanghai) and cloned into pET28-MHL (Genbank accession number: EF456735). Then the plasmid was transformed into E. coli BL21 (DE3) and recombinant protein was over-expressed at 16 °C for 20 hours in the presence of 0.5 mM IPTG. Recombinant SETD3 proteins were purified by fast flow Ni-NTA column (GE Healthcare). N-terminal 6xHis-tag of recombinant proteins were removed by Tobacco Etch Virus (TEV) protease. Gel filtration and ion-exchange were employed for further purification. Purified SETD3 proteins, with their purities exceeding 90% as judged by SDS-PAGE, were concentrated to 15 mg/ml and stored at −80°C before further use. The SETD3 mutants were constructed by conventional PCR using the MutanBEST kit (TaKaRa) and further verified by DNA sequencing. The SETD3 mutants were expressed and purified in the same way as the wild type protein.

For kinetic studies, the enzyme and its mutant forms were produced as fusion proteins with the N-terminal 6xHis-tag and purified by Ni-NTA affinity chromatography. Briefly, the *E. coli* cell paste was resuspended in 11 ml lysis buffer consisting of 25 mM Hepes pH 7.5, 300 mM NaCl, 10 mM KCl, 1 mM DTT, 2 mM MgCl_2_, 1 mM PMSF, 0.25 mg/ml hen egg white lysozyme (BioShop, Canada) and 250 U Viscolase (A and A Biotechnology, Poland). The cells were lysed by freezing in liquid nitrogen and, after thawing and vortexing, the extracts were centrifuged at 4° C (20000 x g for 20 min). For the purification, the supernatant of *E. coli* lysate (11 ml) was diluted 3-fold with buffer A (50 mM Tris-HCl pH 7.4, 400 mM NaCl, 10 mM KCl and 1 mM DTT) and applied onto a HisTrap FF crude column (1 ml) equilibrated with the 90% buffer A and 10% buffer B (30 mM imidazole). The column was then washed with 12 ml 90% buffer A and 10% buffer B, and the retained proteins were eluted with a stepwise gradient of imidazole (5 ml of 60 mM, 5 ml of 150 mM and 5 ml of 300 mM) in buffer A. The recombinant proteins eluted at the highest concentration of imidazole was further processed. The enzyme preparation was desalted onto PD-10 columns equilibrated with 20 mM Tris-HCl pH 7.2, 50 mM KCl, 1 mM DTT and 6% sucrose. The yield of recombinant proteins ranged from 1.6 mg to 0.1 mg for the wild-type and N278A mutant, respectively, per 200 ml of culture. The purified proteins were aliquoted and stored at –70°C.

### Overexpression and purification of the recombinant β-actin inclusion-body protein

Recombinant human β-actin (ACTB, GenBank: NM_001101.4) was prepared as described before (Kwiatkowski et al., 2018). Plasmid pCOLD I that encodes human protein was a kind gift of Dr. Minoru Tamura (Ehime University, Japan) and was prepared as described by (Tamura, 2018).

For β-actin production, Escherichia coli BL21(DE3) (Agilent, USA) cells were transformed with the DNA construct and cultured in 500 mL of LB broth (with 100 μg/ml ampicilin) at 37 °C and 200 rpm until an OD_600_ of 0.5 was reached. Protein production was induced by cold-shock (20 min on ice) and IPTG addition to a final concentration of 0.2 mM. β-actin production was carried out for 20 h at 15 °C, 200 rpm, and harvested by centrifugation (6000 × g for 10 min). The cell paste was resuspended in 27.5 ml lysis buffer consisted of 20 mM Hepes, pH 7.5, 1 mM DTT, 1 mM ADP, 0.5 mM PMSF, 2 μg/ml leupeptin and 2 μg/ml antipain, 0.2 mg/ml hen egg white lysozyme (BioShop), 1000 U Viscolase (A&A Biotechnology, Poland). The cells were lysed by freezing in liquid nitrogen and after thawing and vortexing, the extracts were centrifuged at 4° C (20,000 × g for 30 min).

The pellet, containing inclusion bodies, was completely resuspended in buffer A (20 mM Hepes, pH 7.5, 2 M urea, 0.5 M NaCl, 5 mM DTT, 2 mM EDTA) with the use of Potter-Elvehjem homogenizer and centrifuged at 4° C (20,000 × g for 10 min). The resulting pellet was then subjected to a further two rounds of sequential wash in buffer B (20 mM Hepes, pH 7.5, 0.5 M NaCl, 5 mM DTT, 2 mM EDTA) and buffer C (20 mM Hepes, pH 7.5, 0.5 M NaCl). The washed inclusion bodies were finally solubilized in loading buffer (20 mM Tris-HCl, pH 7.5, 6 M guanidine HCl, 0.5 M NaCl, 10 mM imidazole) and applied on a HisTrap FF column (5 ml) equilibrated with the same buffer.

The column was washed with 20 ml of loading buffer and the bound protein was refolded by the use of a linear gradient of 6-0 M guanidine HCl in loading buffer (40 ml for 20 min). Next, the column was washed with 15 ml of loading buffer without guanidine HCl and the retained proteins were eluted with a stepwise gradient of imidazole (25 ml of 40 mM, 25 ml of 60 mM and 20 ml of 500 mM). The recombinant proteins were eluted with 500 mM imidazole in homogeneous form as confirmed by SDS-PAGE (not shown). The β-actin preparation (10 ml) was immediately dialyzed against 400 ml of the buffer consisted of 20 mM Tris-HCl, pH 7.5, 1 mM DTT, 6% sucrose, 2 μg/ml leupeptin and 2 μg/ml antipain. The dialysis buffer was exchanged three times with the following dialysis time: 2, 2 and 12 h. The purified β-actin was stored at −70°C.

### Overexpression and purification of the recombinant SAH nucleosidase and adenine deaminase

*E. coli* DNA was extracted by heating 50 ml of over-night cultured BL21(DE3) cells at 95°C for 15 min, whereas *Bacillus subtilis* (ATCC 6633, Sigma-Aldrich) genomic DNA was purified from 100 mg of bacterial cells with the use of TriPure reagent according to the manufacturer’s instructions.

The open reading frames encoding *E. coli* SAH nucleosidase and *B. subtilis* adenine deaminase (NCBI Reference Sequence: NC_000913.3 and NC_000964.3, respectively) were PCR-amplified using either Pfu DNA polymerase alone or a mixture of Taq:Pfu polymerases (1:0.2), respectively, in the presence of 1 M betaine. SAH nucleosidase ORF was amplified using a 50 primer containing the initiator codon preceded by an NdeI site and a 30 primer with a HindIII site, whereas adenine deaminase DNA was amplified using a 50 primer with the initiator codon preceded by an KpnI site and a 30 primer with a BamHI site (for primer sequences, see Table 5). The amplified DNA products of expected size were digested with the appropriate restriction enzymes, cloned into the pCOLD I expression vector (pCOLD I/SAH nucleosidase and pCOLD I/adenine deaminase) and verified by DNA sequencing (Macrogen, The Netherlands).

For protein production, *E. coli* BL21(DE3) cells were transformed with the appropriate DNA construct and a single colony was selected to start an over-night pre-culture. 100 mL of LB broth (with 100 mg/mL ampicilin) was inoculated with 10 ml of the pre-culture and incubated at 37°C and 200 rpm until an OD600 of 0.6 was reached. The culture was placed on ice for 20 min (cold-shock) and IPTG was added to a final concentration of 0.25 mM to induce protein expression. Cells were incubated for 16 hr at 13°C, 200 rpm, and harvested by centrifugation (6000 x g for 10 min). The cel pellet was resuspended in 10 ml lysis buffer consisting of 25 mM Hepes pH 7.5, 300 mM NaCl, 50 mM KCl, 1 mM DTT, 2 mM MgCl2, 1 mM PMSF, 5 mg/ml leupeptin and 5 mg/ml antipain, together with 0.25 mg/ml hen egg white lysozyme (BioShop) and 25 U DNase I (Roche). The cells were lysed by freezing in liquid nitrogen and, after thawing and vortexing, the extracts were centrifuged at 4°C (20,000 x g for 30 min).

For protein purification, the supernatant of the *E. coli* lysate (10 ml) was diluted 3-fold with buffer A (50 mM Tris-HCl, pH 7.2, 400 mM NaCl, 10 mM KCl, 30 mM imidazole, 1 mM DTT, 3 mg/ml leupeptin and 3 mg/ml antipain) and applied onto a HisTrap FF column (5 ml) equilibrated with the same buffer. The column was washed with 20–30 ml buffer A and the retained protein was eluted with a stepwise gradient of imidazole (25 ml of 60 mM, 20 ml of 150 mM and 20 ml of 300 mM) in buffer A.

The recombinant proteins were eluted with 150–300 mM imidazole in homogeneous form, as confirmed by SDS-PAGE (not shown). The enzyme preparations were desalted onto PD-10 columns equilibrated with 20 mM Tris-HCl pH 7.2, 50 mM KCl, 1 mM DTT, 6% sucrose, 2 mg/ml leupeptin and 2 mg/ml antipain. The yield of recombinant proteins was 1.2 mg and 3.1 mg of homogenous adenine deaminase and SAH nucleosidase, respectively, per 200 ml of *E. coli* culture. The purified enzymes were aliquoted and stored at –70°C.

### Isothermal titration calorimetry (ITC)

All peptides were synthesized by GL Biochem (Shanghai) Ltd. and were dissolved in water as a stock of 5-12 mM, with the pH of stock solutions adjusted to pH 7.5. Peptides and concentrated proteins were diluted with ITC buffer (20 mM Tris, pH 7.5 and 150 mM NaCl). ITC experiments were performed by titrating 2 μl of peptide (1.2-1.5 mM) into cell containing 50 μM proteins on MicroCal^™^ (Malvern Panalytical, UK) at 25°C, with a spacing time of 160 s and a reference power of 10 μCal/s. Control experiments were performed by injection of peptides into buffer. Binding isotherms were plotted, analyzed and fitted in a one-site binding model by MicroCal PEAQ-ITC Analysis Software (Malvern Panalytical, UK) after subtraction of respective controls. The dissociation constants (Kds) were from a minimum of two experiments (mean ± sd). The ITC binding curves are shown in Supplemental File 1.

### Crystallization, data collection and structure determination

All crystals were grown by using sitting drop vapor diffusion method at 18°C. For crystallization of SETD3 with methylated peptide, SETD3 (12 mg/ml) was pre-incubated with synthesized β-actin peptide (66-88) (GL Biochem Ltd.) and AdoMet at a molar ratio of 1:3:4, and mixed with the crystallization buffer containing 0.1 M sodium cacodylate trihydrate pH 6.5, 0.2M magnesium acetate tetrahydrate, and 20% v/v polyethylene glycol 8000. For the crystallization of SETD3 with unmodified peptide, SETD3, in the concentration of 12 mg/ml, was pre-incubated with synthesized β-actin peptide (66-88) and AdoHcy at a molar ratio of 1:3:4, and mixed with the crystallization buffer containing 0.1M HEPES sodium pH 7.5, 2% v/v polyethylene glycol 400, and 2.0 M ammonium sulfate. Before flash-freezing crystals in liquid nitrogen, all crystals were soaked in a cryo-protectant consisting of 90% reservoir solution plus 10% glycerol. The diffraction data were collected on BL17U1 at the Shanghai Synchrotron Facility (Wang et al., 2016) (SSRF). Data sets were collected at 0.9789 Å or 0.9789 Å, and were processed by using the HKL2000 program (Otwinowski and Minor, 1997).

The initial structures of the SETD3-actin complexes were solved by molecular replacement using Phaser (McCoy et al., 2007) with previously solved SETD3 structure (PDB: 3SMT) as the search model. Then all the models were refined manually and built with Coot (Emsley and Cowtan, 2004). The final structures were further refined by PHENIX (Adams et al., 2010). The statistics for data collection and refinement are summarized in Table 2.

### Mass Spectrometry

Reversed-phase microcapillary/tandem mass spectrometry (LC/MS/MS) was performed using an Easy-nLC nanoflow HPLC (Proxeon Biosciences) with a self-packed 75 μm id x 15 cm C18 column connected to either an QE-Plus (Thermo Scientific) in the data-dependent acquisition and positive ion mode at 300 nL/min. Passing MS/MS spectra were manually inspected to be sure that all b- and y- fragment ions aligned with the assigned sequence and modification sites. A 25-ul reaction mixture containing 2 μM SETD3 or SETD3 mutants (final concentration), 20 μM peptide (final concentration) in a buffer containing 10 mM Tris-HCl, pH 7.5, 20 mM NaCl and 10 μM AdoMet. The reaction was incubated at 37°C for 2 h before being quenched (at 70°C for 10-15 mins). Then reactions were analyzed by LC/MS/MS and Proteomics Browser Software, with the relative abundances of substrate and product reflecting the methylation activities of proteins.

### Radiochemical Assay of the SETD3 Activity

The enzyme activity was determined by measuring the incorporation of [^3^H]methyl group from *S*-[methyl-^3^H]adenosyl-L-methionine ([^3^H]AdoMet) into homogenous recombinant human (mammalian) β-actin or its mutated form in which histidine 73 was replaced by alanine residue (H73A). The standard incubation mixture (0.06-0.11 ml) contained 25 mM Tris-HCl, pH 7.2, 10 mM KCl, 1 mM DTT and various concentration of protein substrate and [^1^H+^3^H] AdoMet (≈ 300-700 × 10^3^ cpm) as indicated in legends to figures and tables. The incubation mixture was supplemented with recombinant AdoHcy nucleosidase and adenine deaminase to prevent AdoHcy accumulation as indicated in legends to figures and tables. The reaction was started by the addition of enzyme preparation and carried out at 37°C for 10 min. Protein methylation was linear for at least 15 min under all conditions studied. By analogy to assays of nonribosomal peptide synthetase activity (Richardt et al., 2003; Drozak et al., 2014), the incubation was stopped by the addition of 0.05-0.11 ml of the reaction mixture to 0.025 ml of BSA (1 mg) and 0.8 ml of ice-cold 10% (w/v) trichloroacetic acid (TCA). After 10 min on ice, the precipitate was pelleted and washed twice with ice-cold 10% TCA. The pellet was finally dissolved in pure formic acid.

### Calculations

V_max_, K_M_ and k_cat_ for the methyltransferase activity of the studied enzymes were calculated with Prism 8.0 (GraphPad Software, La Jolla, USA) using a nonlinear regression.

## ACCESSION NUMBERS

The coordinates and structure factors of AdoHcy bound SETD3 with unmodified β-actin peptide and methylated β-actin peptide haven been deposited into Protein Data Bank (PDB) with accession numbers 6ICV and 6ICT, respectively.

## SUPPLEMENTARY INFORMATION

Supplemental File can be found online at XXXX

## ACKNOWLEDGEMENT

We are grateful to the staff members at beam lines BL17U1, BL18U1 and BL19U1 at Shanghai Synchrotron Radiation Facility for assistance in data collection. This work was supported by National Natural Science Foundation of China Grants 31770806 (to C. X.), 31570737 (to C. X.), 31500601 (to S. L.), and 31501093 (to H. Y.). J.D is supported by Narodowe Centrum Nauki (Opus Grant UMO-2017/27/B/NZ1/00161) (to J.D.). H. Y. is supported by China Postdoctoral Science Foundation Grant (No. 2015M580547). C.X is also supported by the Major/Innovative Program of Development Foundation of Hefei Center for Physical Science and Technology (2018CXFX007) and the “Thousand Young Talent program”. The Structural Genomics Consortium is a registered charity (no. 1097737) that receives funds from AbbVie; Bayer Pharma AG; BoehringerIngelheim; Canada Foundation for Innovation; Eshelman Institute for Innovation; Genome Canada through the Ontario Genomics Institute; the Innovative Medicines Initiative (European Union/European Federation of Pharmaceutical Industries and Associations; Unrestricted Leveraging of Targets for Research Advancement and Drug Discovery [ULTRA-DD] grant no. 115766); Janssen, Merck, and Company; Novartis Pharma AG; Ontario Ministry of Economic Development and Innovation; Pfizer; São Paulo Research Foundation (FAPESP); Takeda; and the Wellcome Trust (to J.M.).

## CONFLICT OF INTEREST

The authors declare that they have no conflict of interest related to the publication of this manuscript.

## FIGURE SUPPLEMENTS

**Suppplemental File 1.** ITC binding curves for the binding measurements reported in Table 1.

## TITLES FOR THE SOURCE DATA FILES

Figure 3 **– Source Data 1.** Radiochemical measurements of SETD3-dependent methylation of either human recombinant β-actin or its mutated form (H73A) in the presence of increasing pH value of the reaction mixture.

Table 3 **– Source Data 1**. Determination of kinetic parameters of SETD3-catalyzed methylation of actin (for β-actin as the substrate).

Table 3 **– Source Data 2.** Determination of kinetic parameters of SETD3-catalyzed methylation of actin (for AdoMet as the substrate).

**Figure 1 - figure supplement 1.**
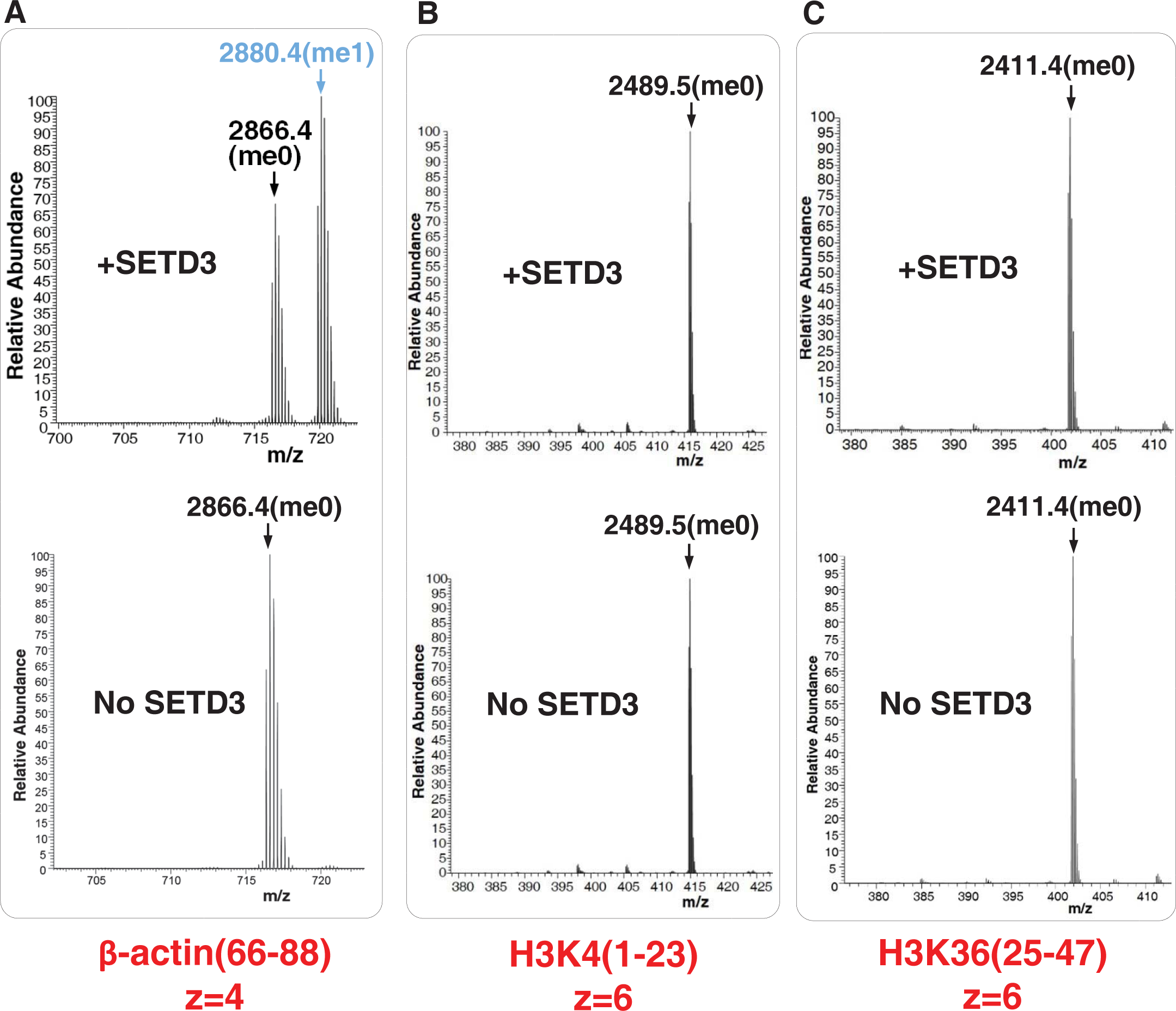
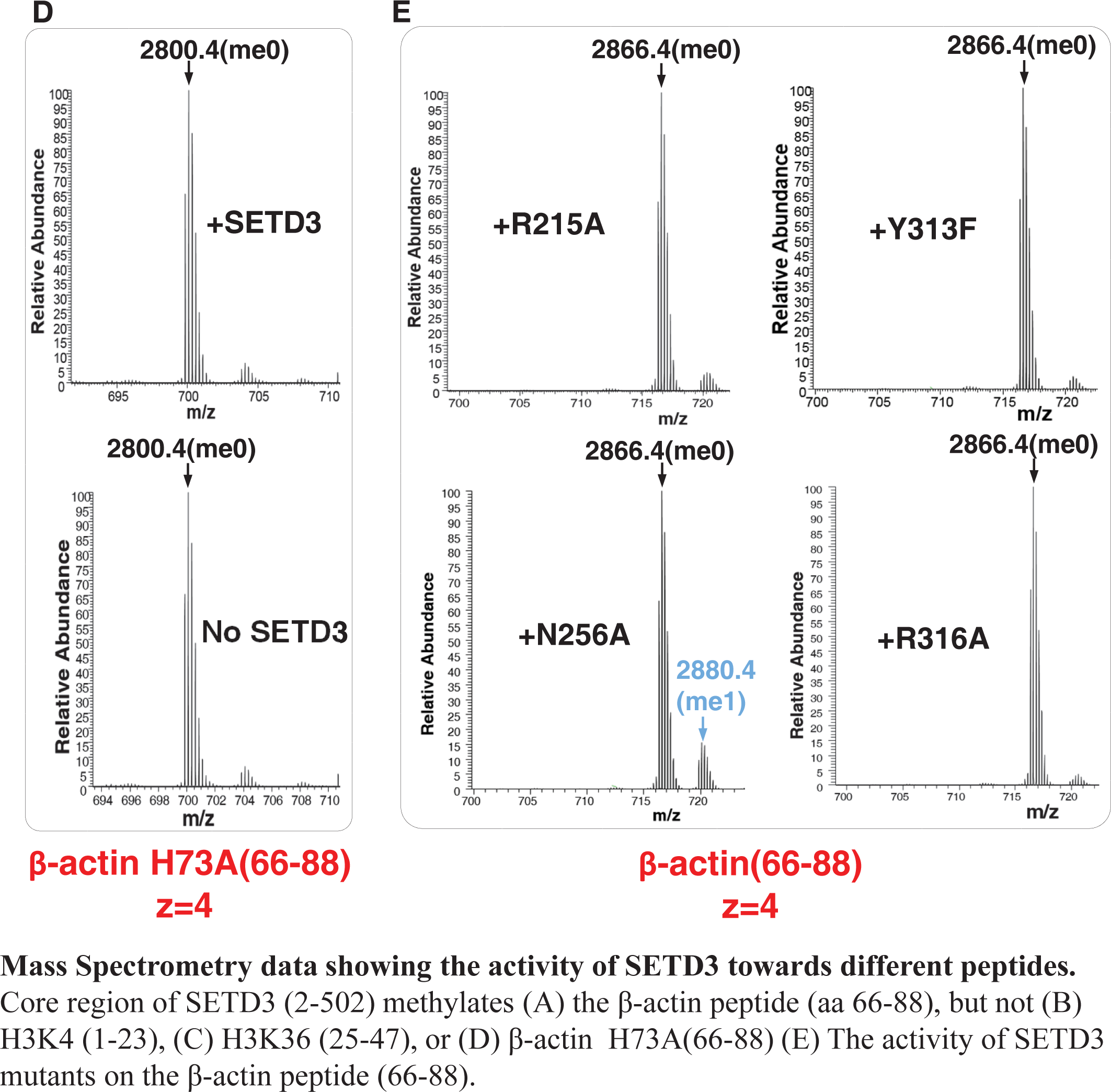
Mass Spectrometry data showing the activity of SETD3 towards different peptides. Core region of SETD3 (2-502) methylates (A) the β-actin peptide (aa 66-88), but not (B) H3K4 (1-23), (C) H3K36 (25-47), or (D) β-actin H73A(66-88) (E) The activity of SETD3 mutants on the β-actin peptide (66-88).

**Figure 1 - figure supplement 2.**
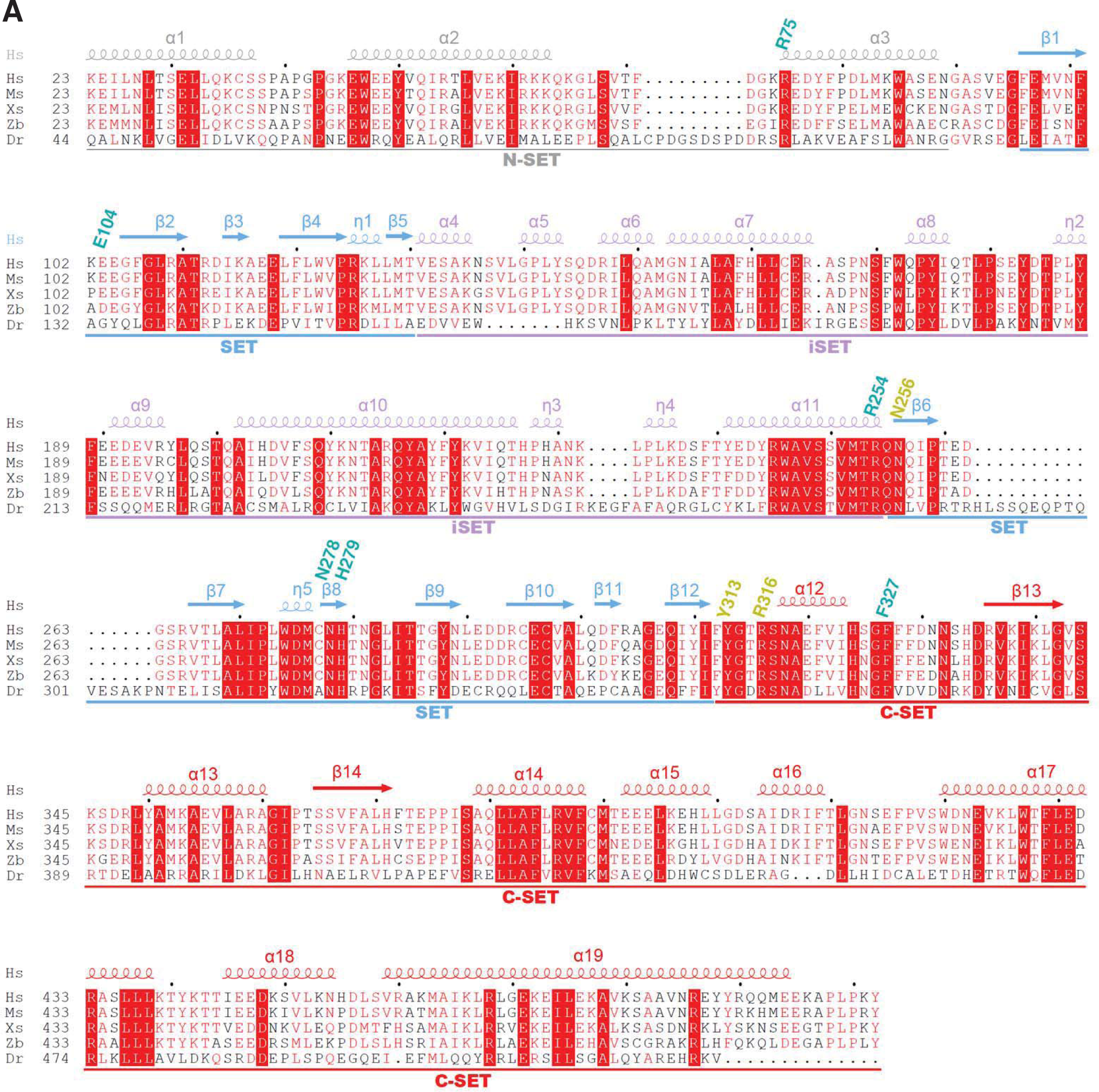
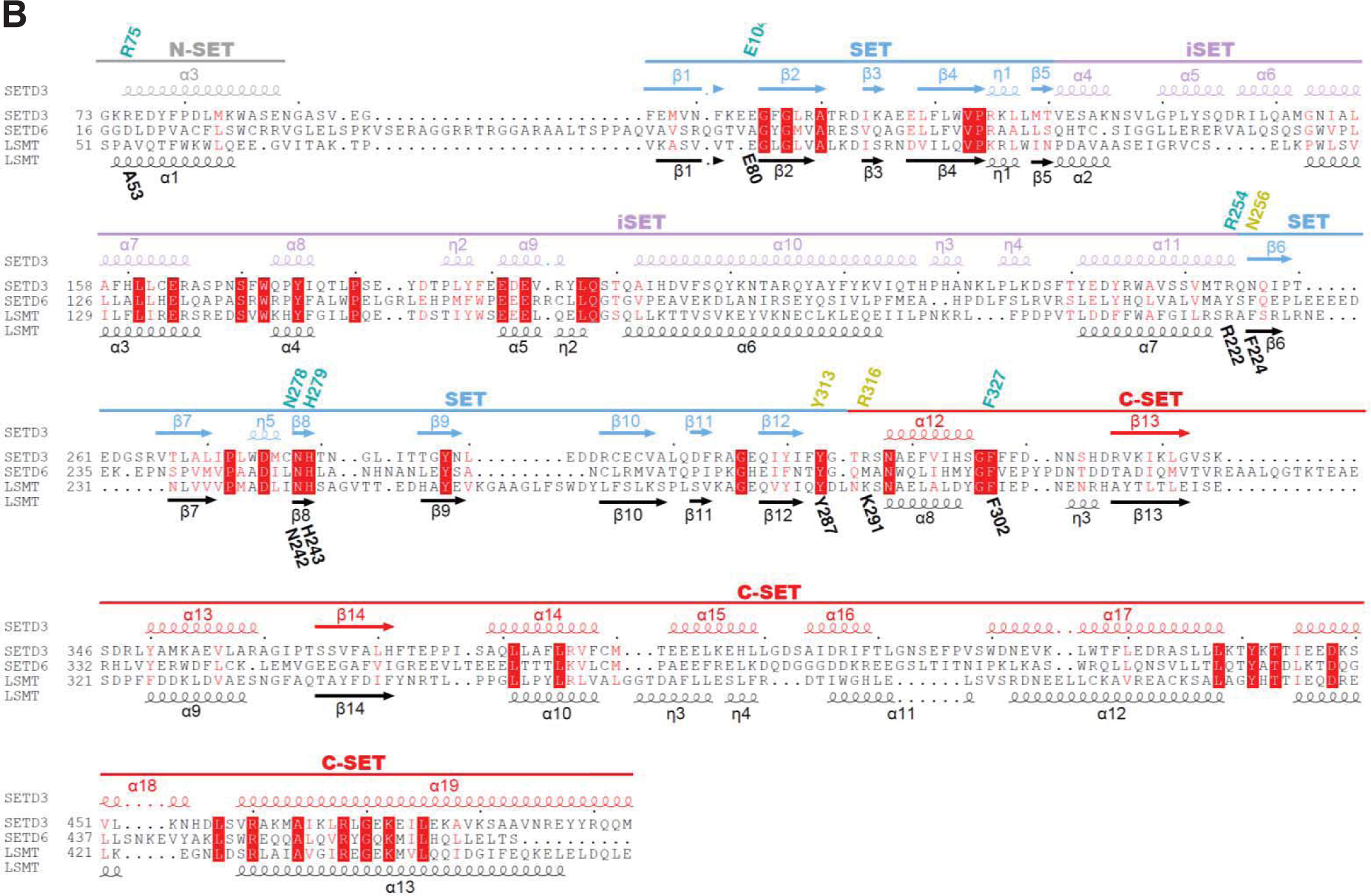
Sequence alignment of human SETD3 with its orthologs or other SET domain proteins (SETD6 and LSMT). (A) Sequence alignment of Human SETD3(Hs, NP_115609.2), mouse SETD3(Ms, NP_082538.2), Xenopus SETD3(Xs, NP_001016577), Zebrafish SETD3 (Zb, NP_956348.1) and Drosophila (Dr, XP_017048262.1). The secondary structures of human SETD3 are colored in the same way as shown in Figure 1A. The residues involved in binding to AdoHcy and His73 are labeled in cyan and yellow, respectively. (B) Sequence aligment of human SETD3, human SETD6 (NP_001153777.1) and Pisum sativum LSMT (AAA69903.1). The secondary structures, AdoHcy binding residues, and His73 binding residues, are labeled in the same way as shown in Figure 1–figure supplement 2A.

**Figure 1 - figure supplement 3.**
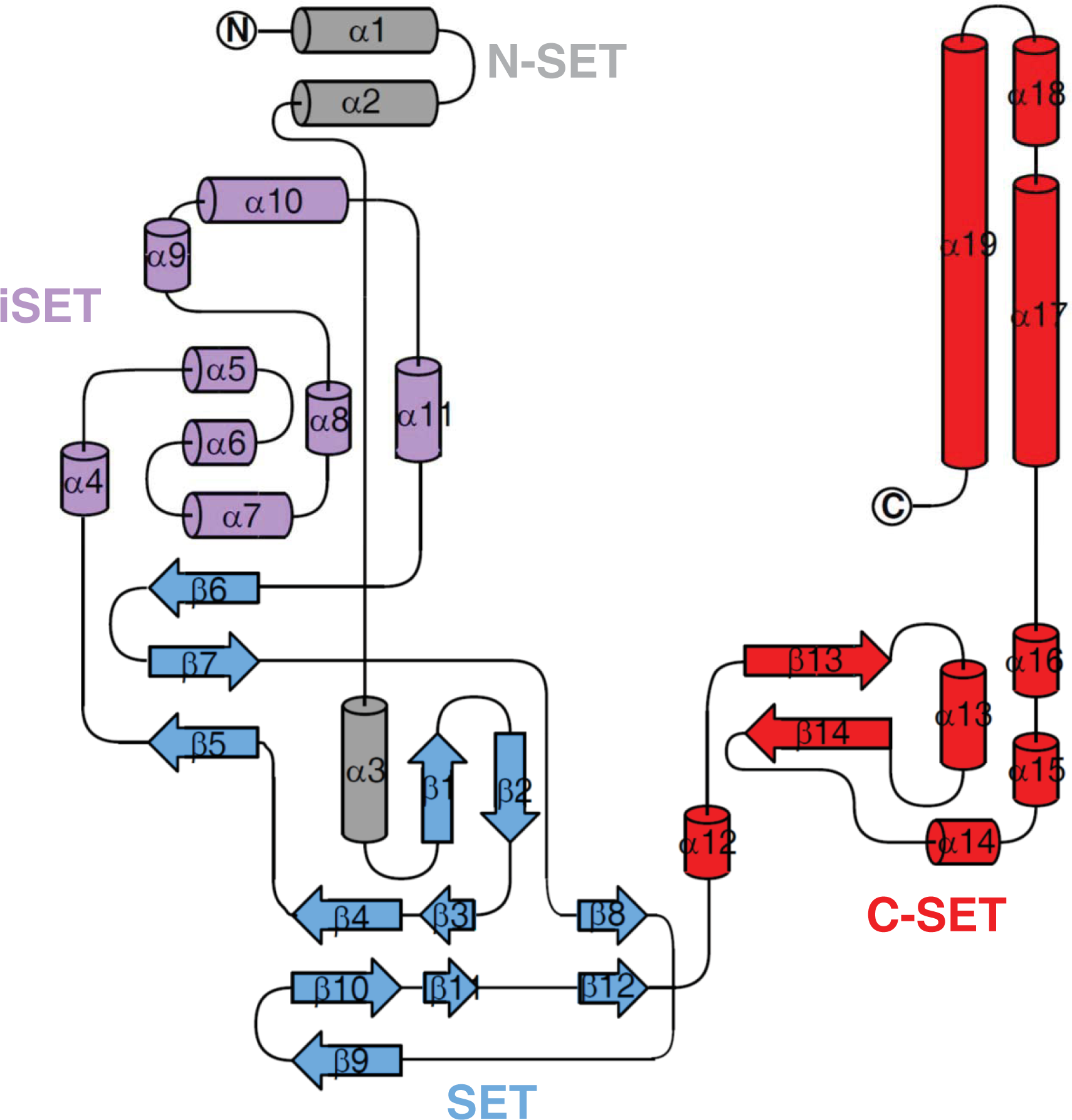
**Topology of SETD3 (2-502)**, with secondary structures marked and colored in the same way as shown in Figure 1–figure supplement 2A.

**Figure 1 - figure supplement 4.**
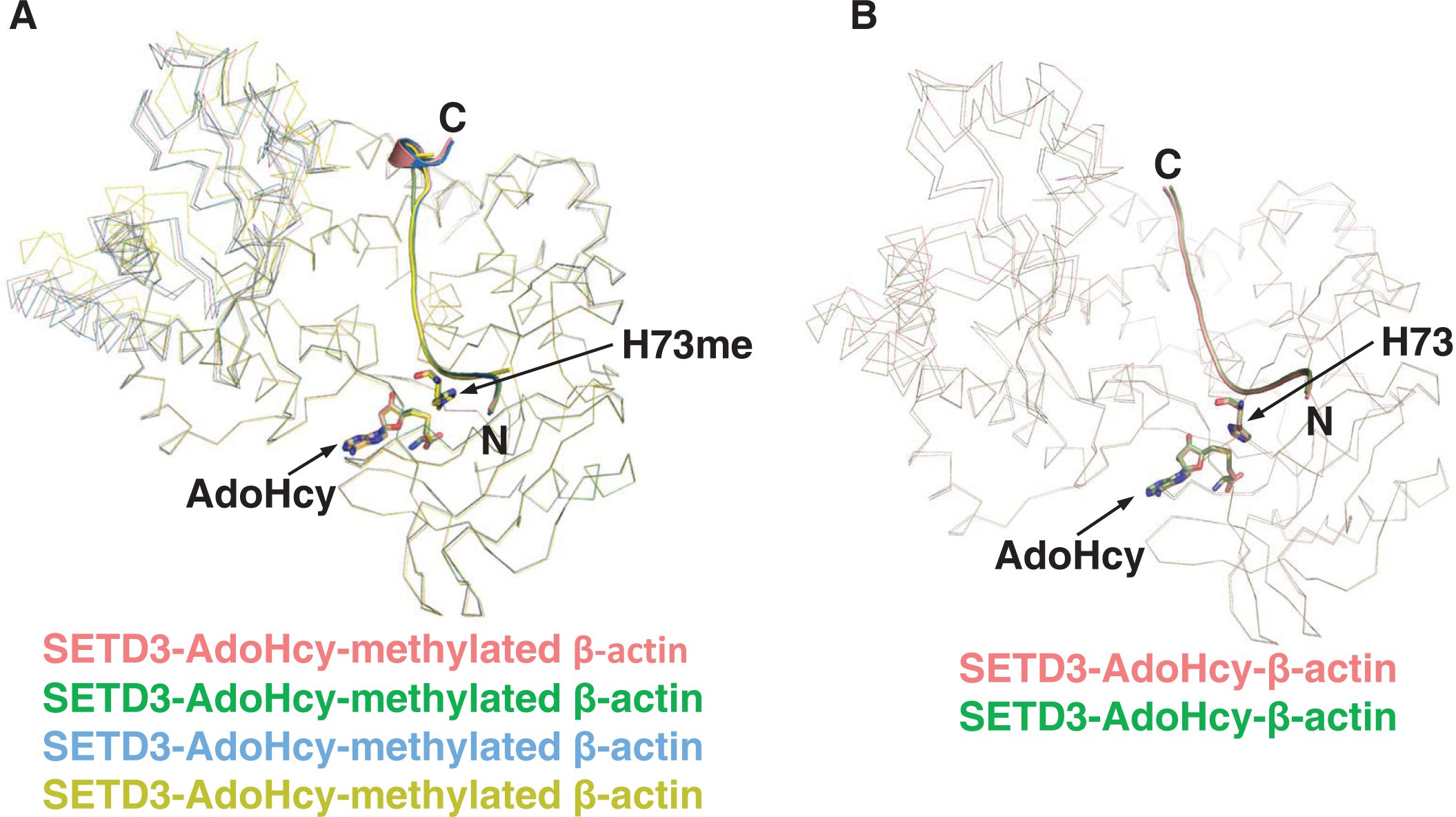
Superposition of SETD3 molecules. (A) Superposition of four SETD3 molecules bound to methylated β-actin in the same asymmetric unit, with the four molecules shown in red, green, blue and yellow ribbon, respectively. His73me of β-actin and AdoHcy are shown in sticks. (B) Superposition of two molecules of SETD3 bound to unmodified β-actin in the same asymmetric unit, with the two molecules shown in red and green ribbon, respectively. His73 of β-actin and AdoHcy are shown in sticks.

**Figure 1 - figure supplement 5.**
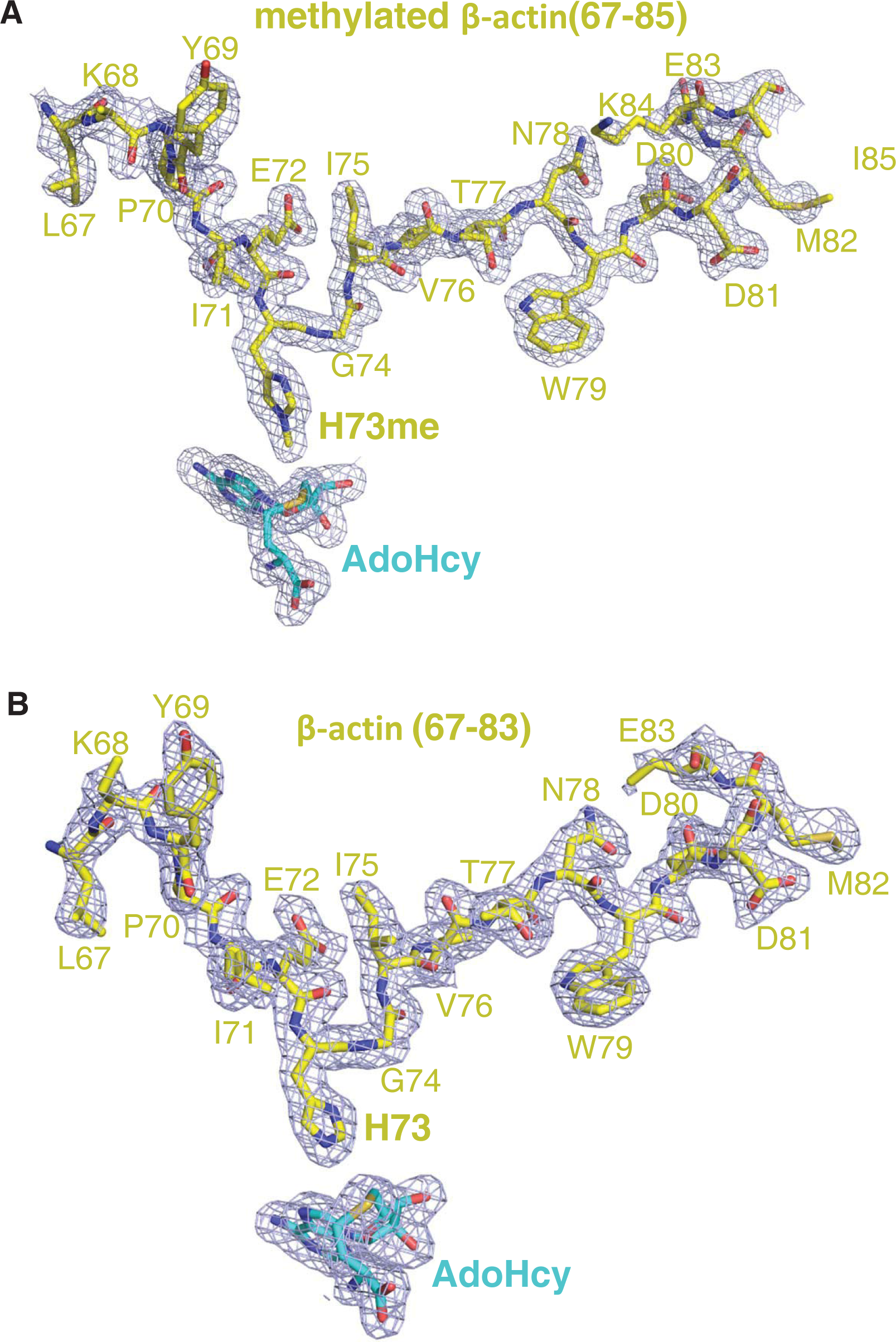
**The 2|Fo|–|Fc| σ-weighted maps of (A) methylated β-actin peptide and (B) unmodified β-actin peptide in the presence of AdoHcy**, are contoured at 1.2 σ (blue cage), respectively. The peptides are shown in yellow sticks.

**Figure 1 - figure supplement 6.**
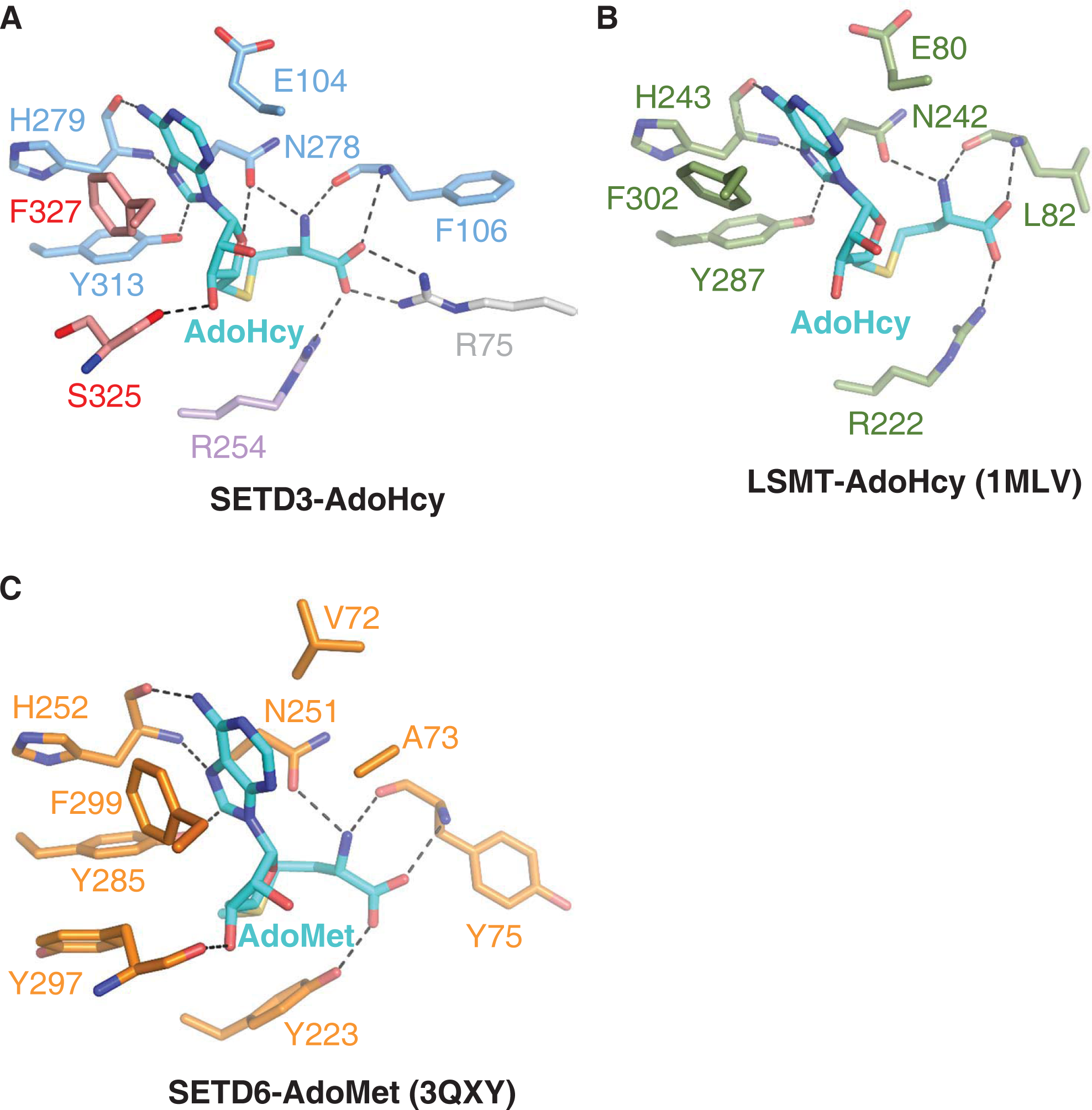
Comparison of AdoHcy binding mode of SETD3 with those observed in LSMT and SETD6. (A) A detailed depiction of intermolecular interactions between SETD3 and AdoHcy, with AdoHcy shown in cyan sticks, and the AdoHcy binding residues of SETD3 shown in sticks and colored in the same mode as in Figure 1A. (B) Detailed interactions between AdoHcy and LSMT (PDB id: 1MLV), with AdoHcy binding residues of LSMT and AdoHcy shown in green and cyan sticks, respectively. (C) Detailed interactions between AdoMet and SETD6 (PDB id: 3QXY), with AdoMet binding residues of SETD6 and AdoMet shown in orange and cyan sticks, respectively. In Figures 2B and 2C, hydrogen bonds are indicated by black dashes.

**Figure 1 - figure supplement 7.**
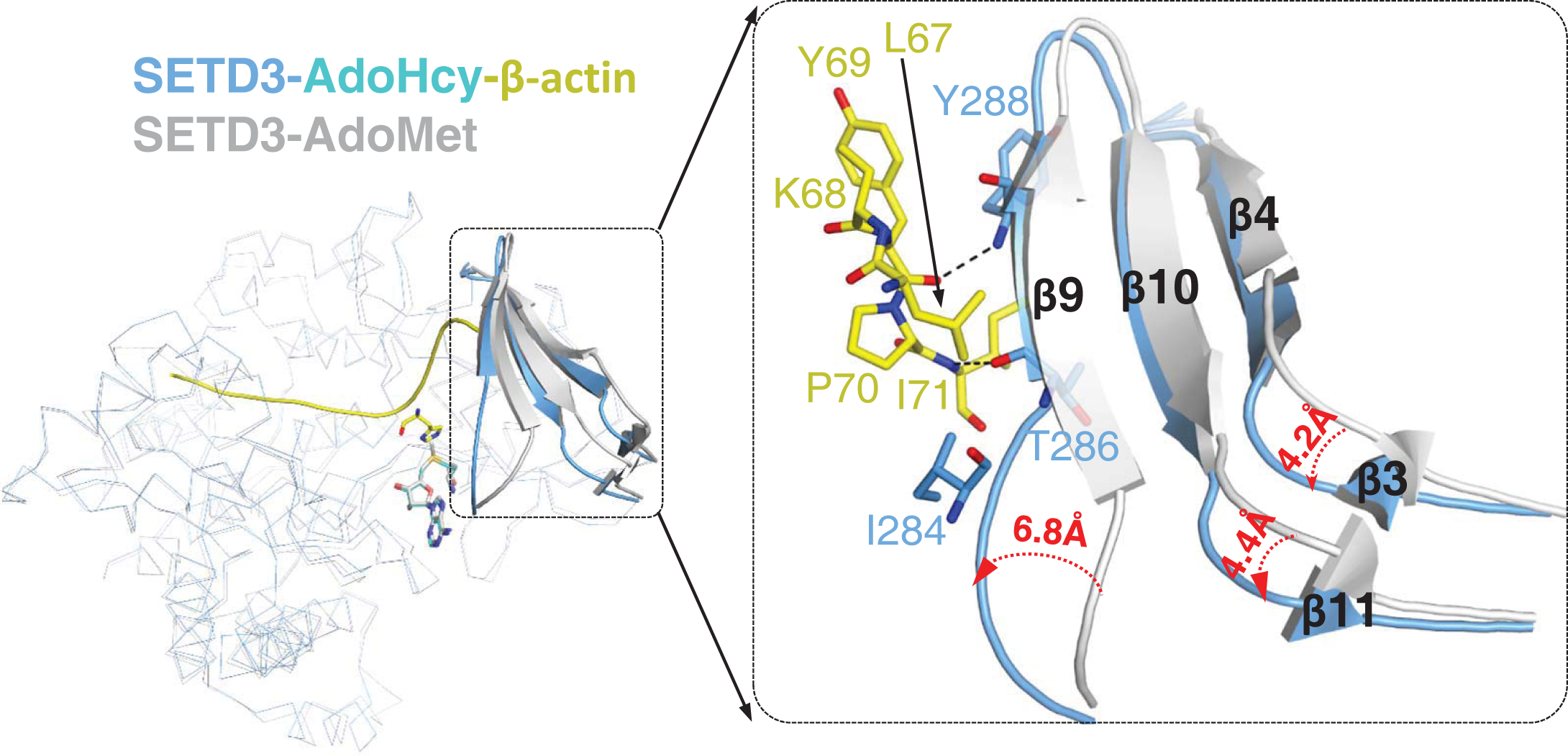
Superposition of AdoMet-bound SETD3 (PDB id: 3SMT) with AdoHcy bound SETD3-actin. In the structure of AdoMet bound SETD3, the protein is shown in grey ribbon and AdoMet is shown in grey sticks. In the structure of AdeHcy bound SETD3 in the presence of actin, protein is shown in blue ribbon and AdoHcy is shown in yellow sticks. β4-β9-β10 and β3-β11 changed conformations significantly upon binding to actin peptide, which are shown in cartoon representation. Leu67-Ile71 of β-actin and their interaction residues in SETD3, are shown in sticks. Three loops that precede β4, β9 and β11 shift 4.2 Å, 4.4 Å, and 6.8 Å, respectively.

**Figure 1 - figure supplement 8.**
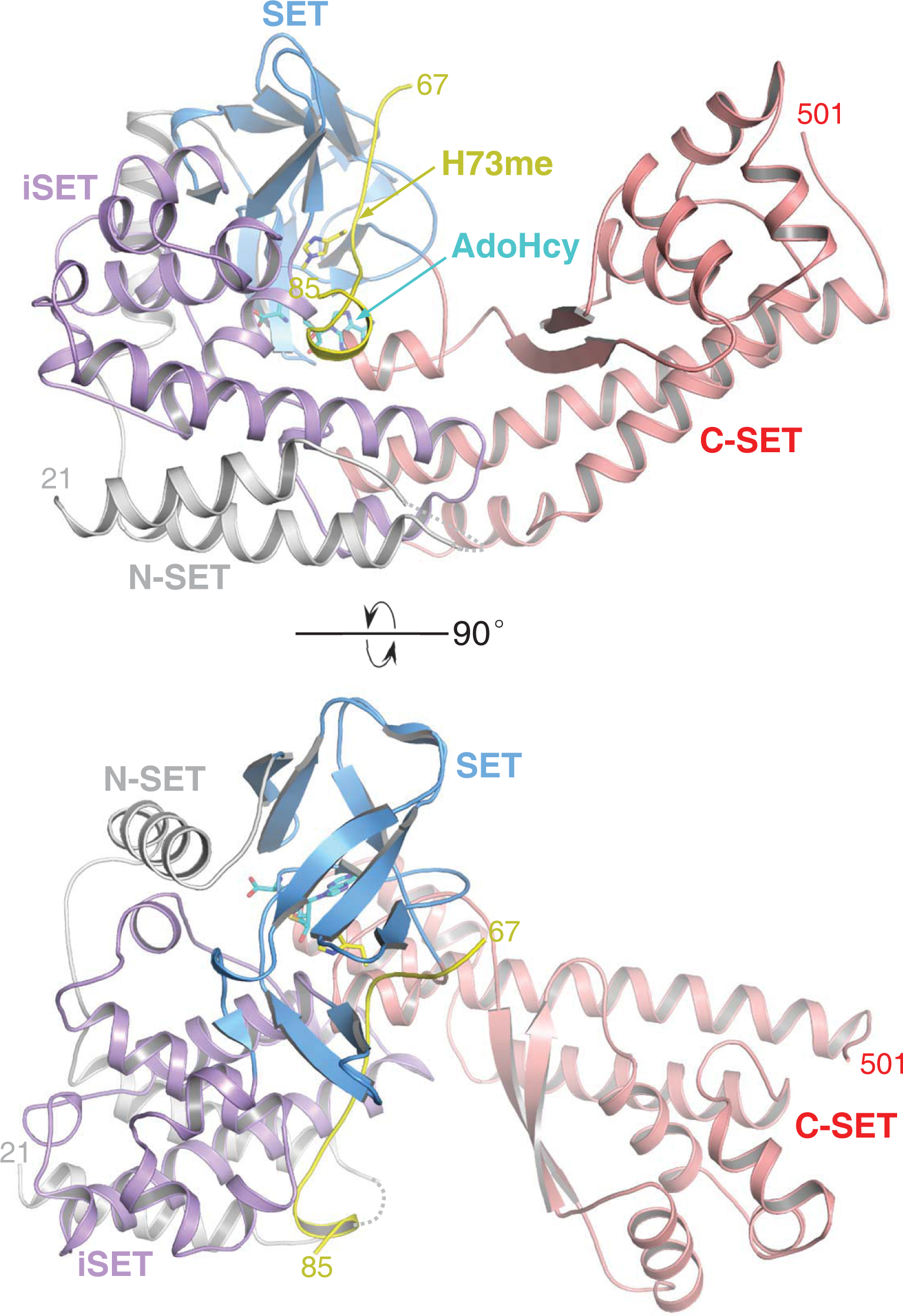
Overall structure of SETD3 with methylated β-actin peptide in the presence of AdoHcy. SETD3 is colored and labelled in the same mode as shown in Figure 1C. AdoHcy and His73me are shown in sticks.

**Figure 2 - figure supplement 1.**
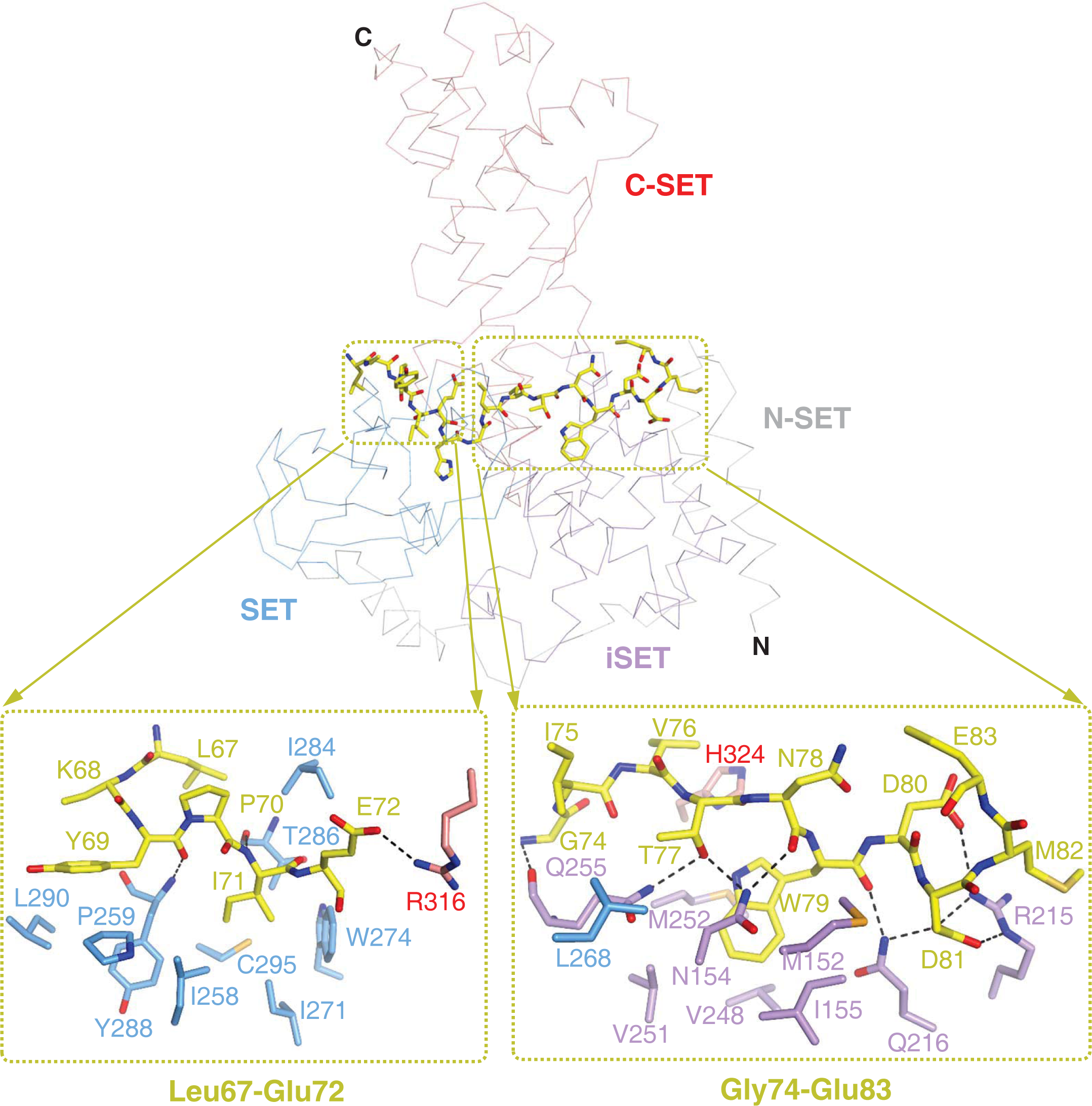
Detailed interactions of SETD3 and β-actin peptide (66-88). The β-actin peptide mainly contacts N-terminal lobe of SETD3, with its N-terminal and C-terminal sides contact SET and iSET domains, respectively. In the top panel, SETD3 are shown in ribbon and colored in the same way as shown in Figure 2A. β-actin peptide is shown in yellow sticks. Detailed interactions between SETD3 and Leu67-Glu72, and between SETD3 and Gly74 and Glu83, are shown in the left bottom and right bottom panels, respectively. The SETD3 residues involved in β-actin binding are shown in sticks.

**Figure 2 - figure supplement 2.**
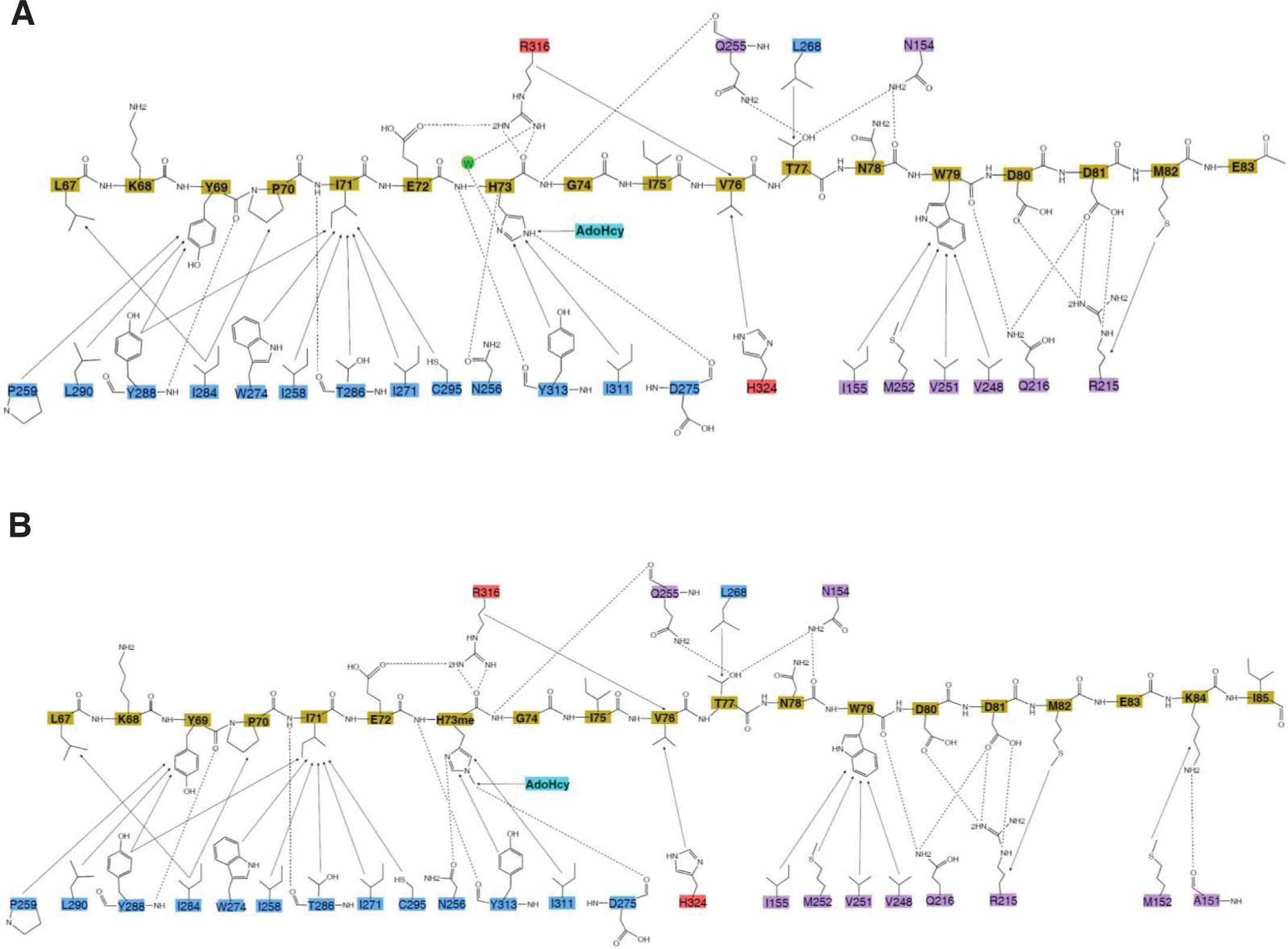
Schematic of the detailed interactions (A) between SETD3 and unmodified β-actin (B) and between SETD3 and methylated β-actin. SETD3 residues are colored as shown in Figure 1A, while the peptide residues are colored in yellow. The hydrophobic interactions and hydrogen bonds between protein and peptide are shown in black solid arrows and black dash arrows, respectively.

**Figure 2 - figure supplement 3.**
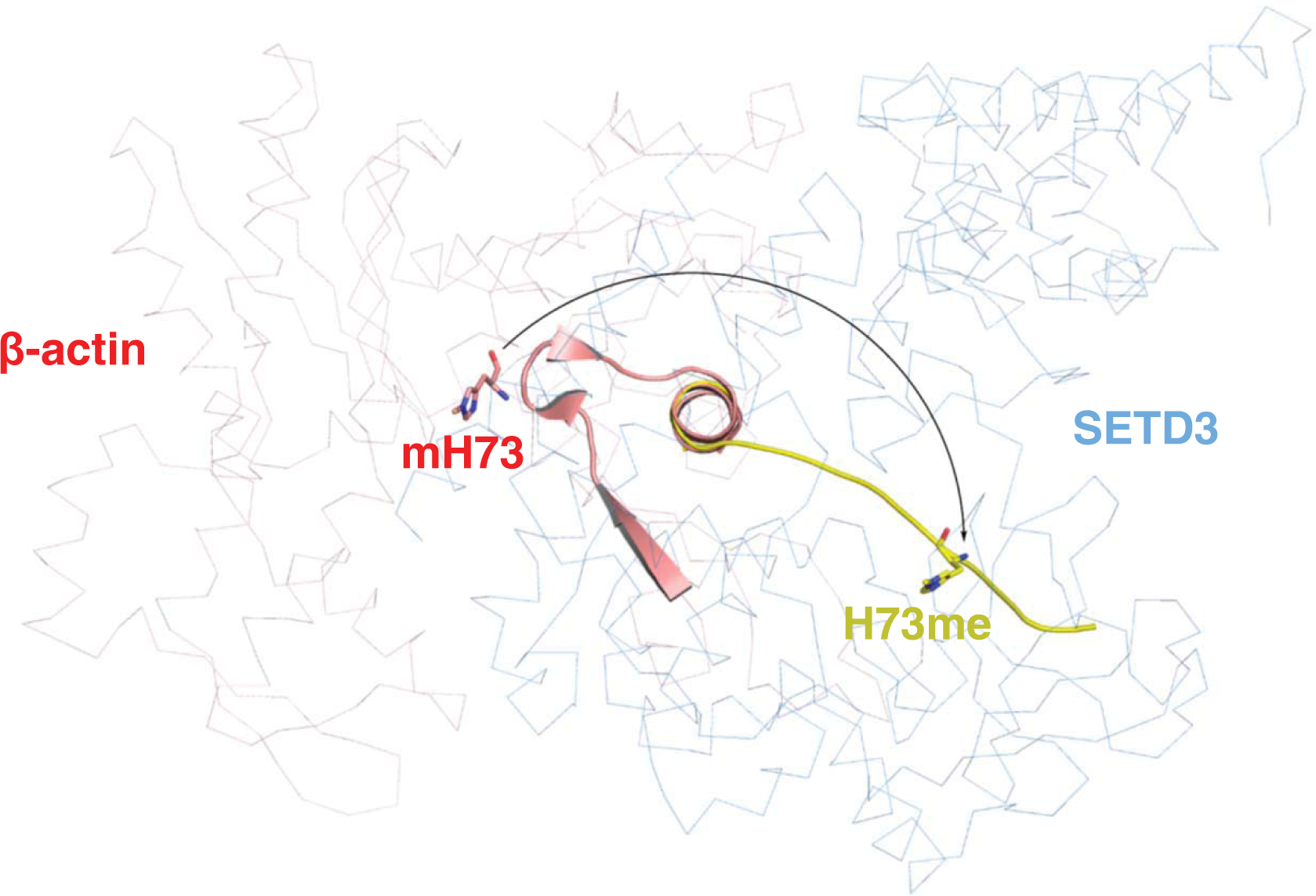
Actin endures conformational changes upon binding to SETD3. β-actin peptide (66-88, yellow cartoon) is over laid with the same fragment in the structure of native β-actin (red cartoon) (PDB id: 1HLU). native β-actin decomposes its local secondary structures upon SETD3 binding. Native β-actin and SETD3 are shown in red and blue ribbon, respectively.

